# Chromatin and transcriptional adaptation accompany monocyte engraftment in the injured retina

**DOI:** 10.1101/2024.09.09.612126

**Authors:** Jie Liu, Fengyang Lei, Chiara Romano, Bin Yan, Tian Cao, Naiwen Cui, Jyoti Sharma, Victor Correa, Lara Roach, Savvas Nicolaou, Kristen Pitts, James Chodosh, Daniel E. Maidana, Demetrios Vavvas, Milica A Margeta, Huidan Zhang, David Weitz, Raul Mostoslavsky, Eleftherios I. Paschalis

## Abstract

Distinguishing resident microglia from engrafted peripheral myeloid cells is important for interpreting CNS inflammation. P2RY12, TMEM119, and FCRLS are widely used to identify microglia, but their ability to distinguish cellular origin after prolonged engraftment remains context dependent. Using bone marrow chimeras, single-cell and bulk RNA sequencing, ATAC-seq, flow cytometry, and immunostaining, we examined myeloid adaptation after retinal injury. Donor-derived cells acquired P2RY12, TMEM119, and FCRLS expression with different kinetics and degrees of heterogeneity, while subsets downregulated CCR2, Ly6C, and CD45. These phenotypic changes were accompanied by altered chromatin accessibility at microglia-associated loci and changes in transcription factor motif enrichment. At one year after injury, engrafted cells and injured host microglia share inflammatory transcriptional features while retaining population-associated differences with significantly lower expression of *Sall1* in engrafted cells compared to naive microglia. Donor-derived cells infiltrating the brain and spinal cord without additional CNS injury also acquired microglial markers. These observations establish phenotypic overlap and persistent molecular differences in engrafted retinal monocytes and support combining marker expression with cellular-origin information to identify CNS myeloid populations after long-term engraftment.

**Significance Statement:** Peripheral CX3CR1 cell infiltration into the retina occurs in response to injury, while the brain and spinal cord become repopulated over time without CNS injury. Nevertheless, engrafted myeloid cells acquired commonly used microglial markers and, in the retina, showed accompanying changes in chromatin accessibility and gene expression. Long-term engrafted cells retained transcriptional differences from host microglia with significantly reduced *Sall1* expression relative to naive microglia. These findings define the extent and persistence of retinal myeloid adaptation and support combining marker profiles with cellular-origin information to distinguish engrafted cells from resident microglia.

## Introduction

Microglia and infiltrating peripheral monocytes contribute to CNS homeostasis and pathology and have emerged as therapeutic targets [1–5]. Distinguishing these populations becomes increasingly challenging after prolonged tissue residence, as engrafted monocytes acquire features that resemble those of resident microglia. Our previous work showed that peripheral monocytes engraft in the injured retina and remain proinflammatory over the long term, with elevated expression of MHC-II, IL-1β, and TNF-α despite adopting a ramified, microglia-like morphology [2]. This persistent inflammatory phenotype underscores the importance of identifying cellular origin when assessing the contributions of myeloid populations to chronic retinal inflammation and developing targeted therapeutic strategies.

P2RY12, TMEM119, and FCRLS have been widely adopted to identify microglia, with their specificity established through comparisons of resident microglia, peripheral monocytes, and selected inflammatory conditions [6, 7]. However, prolonged CNS residence can alter the features that initially distinguish these populations. Monocyte-derived cells can acquire selected microglial markers while retaining molecular or functional differences from resident microglia [2, 8–12]. Marker expression may therefore reflect adaptation to the tissue environment as well as cellular identity. Defining how these phenotypic changes relate to chromatin accessibility and transcription is important for understanding the extent of this adaptation and interpreting myeloid populations after injury and transplantation.

Here we investigated the temporal acquisition of microglial markers by engrafted peripheral myeloid cells and the accompanying changes in chromatin accessibility and transcription. Our principal model was retinal injury in busulfan-conditioned bone marrow chimeras, complemented by observations in the brain and spinal cord of chimeras without additional CNS injury. Donor-derived cells acquired P2RY12, TMEM119, and FCRLS with distinct kinetics, while subsets downregulated CCR2, Ly6C, and CD45.

Chromatin-accessibility profiling and bulk RNA-seq revealed shared features alongside persistent differences relative to host microglia. These findings characterize the molecular adaptation accompanying engraftment and highlight the need to integrate cellular origin with phenotypic and molecular profiles when identifying the myeloid populations involved in long-term retinal inflammation.

## Results

### Single-cell RNA sequencing identifies temporal changes in retinal myeloid transcriptional states

We performed single-cell RNA sequencing (scRNA-seq) on CD45^+^CD11b^+^ cells isolated from mouse retinas before and at 1, 4, and 7 days after corneal alkali injury, a model associated with monocyte infiltration and engraftment [2, 13, 14]. Four transcriptional clusters were identified **(Fig. 1A)**. The baseline population had a resident microglial profile, whereas the day 1 population was more heterogeneous and included cells with increased *Siglec1* expression **(Fig. 1B)** [15]. Later samples showed increased representation of transcriptional states expressing microglia-associated genes.

**Figure 1.**
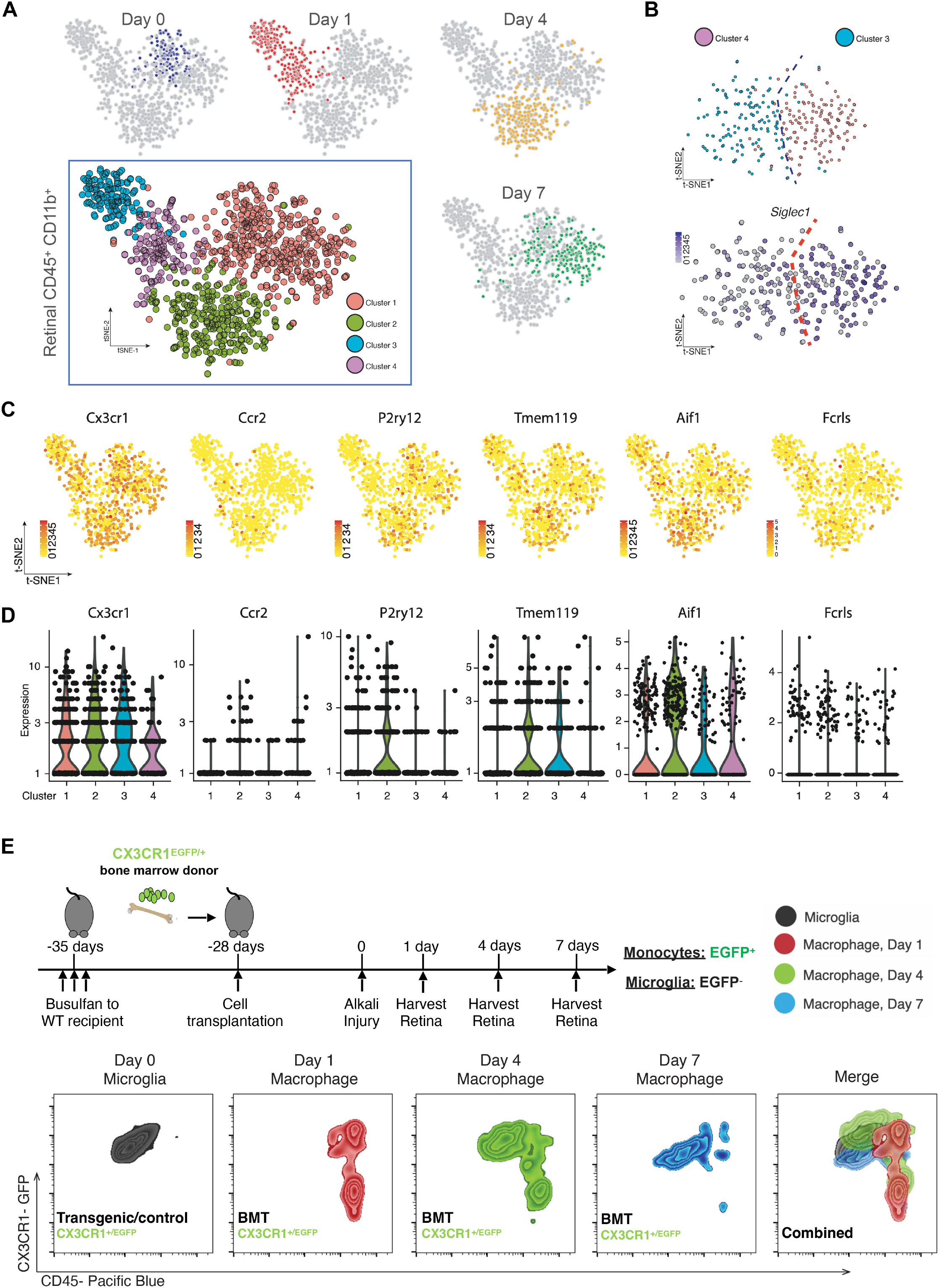
Single-cell transcriptional profiles and CD45 expression after retinal injury. **(A)** t-SNE representation of retinal CD45^+^CD11b^+^ cells collected before and after ocular injury, showing four transcriptional clusters and their distributions across sampling times. **(B)** *Siglec1* expression in selected early post-injury populations. **(C)** Feature plots for *Cx3cr1*, *Ccr2*, *P2ry12*, *Tmem119*, *Aif1*, and *Fcrls*. **(D)** Expression distributions for the six genes shown in C across the four clusters. **(E)** CD45 expression in CX3CR1^+^ retinal cells. An uninjured CX3CR1^+/EGFP^ transgenic mouse provides a reference for the resident-cell profile; CX3CR1^+/EGFP^ bone marrow chimeras show donor-derived cells at the indicated post-injury times. CD45 expression shifts downward in a subset of donor-derived cells during the first week, with overlap with the host microglial range.

*Cx3cr1* was broadly expressed, whereas *Ccr2* expression was enriched in early post-injury populations **(Fig. 1C, D)**. *Tmem119* and *Fcrls* were detected across multiple clusters. *P2ry12* was low or undetectable in the early day 1 clusters and more prominent in the later samples; *Aif1*, which encodes IBA1, was expressed across clusters **(Fig. 1C, D)**. In a complementary CX3CR1+/EGFP bone marrow chimera model [2, 4, 16], flow cytometry showed a shift toward lower CD45 expression in donor-derived retinal cells during the first week after injury, with overlap with the resident microglial range **(Fig. 1E)**. Blood chimerism was approximately 95% in CD45^+^CD11b^+^CX3CR1^+^ cells **(Fig. S1)**. Uninjured CX3CR1^+/EGFP^ transgenic mice provided a reference for the resident microglial CD45 profile.

We assessed selected transcripts by qPCR in sorted cells from bone marrow chimeras 45 days after injury. Engrafted monocytes expressed *P2ry12*, *Tmem119*, *Fcrls*, and *Aif1* transcripts **(Fig. S2)**. *P2ry12* abundance was higher than in blood monocytes but remained below that in naive or injured host microglia.

### Donor-derived retinal cells acquire P2RY12, TMEM119, and FCRLS protein expression

Using CX3CR1^+/EGFP^::CCR2^+/RFP^ bone marrow chimeras to distinguish donor-derived EGFP^+^ cells from host EGFP^−^ cells, we assessed microglial marker expression after retinal injury **(Fig. 2A)**. In the circulating cells examined, immunostaining did not detect P2RY12, TMEM119, or IBA1, whereas MHC-II staining was detectable **(Fig. S3)**. Flow cytometry also showed low or undetectable FCRLS in CD45^+^CD11b^+^CX3CR1^+^ blood cells **(Fig. 2D)**.

**Figure 2.**
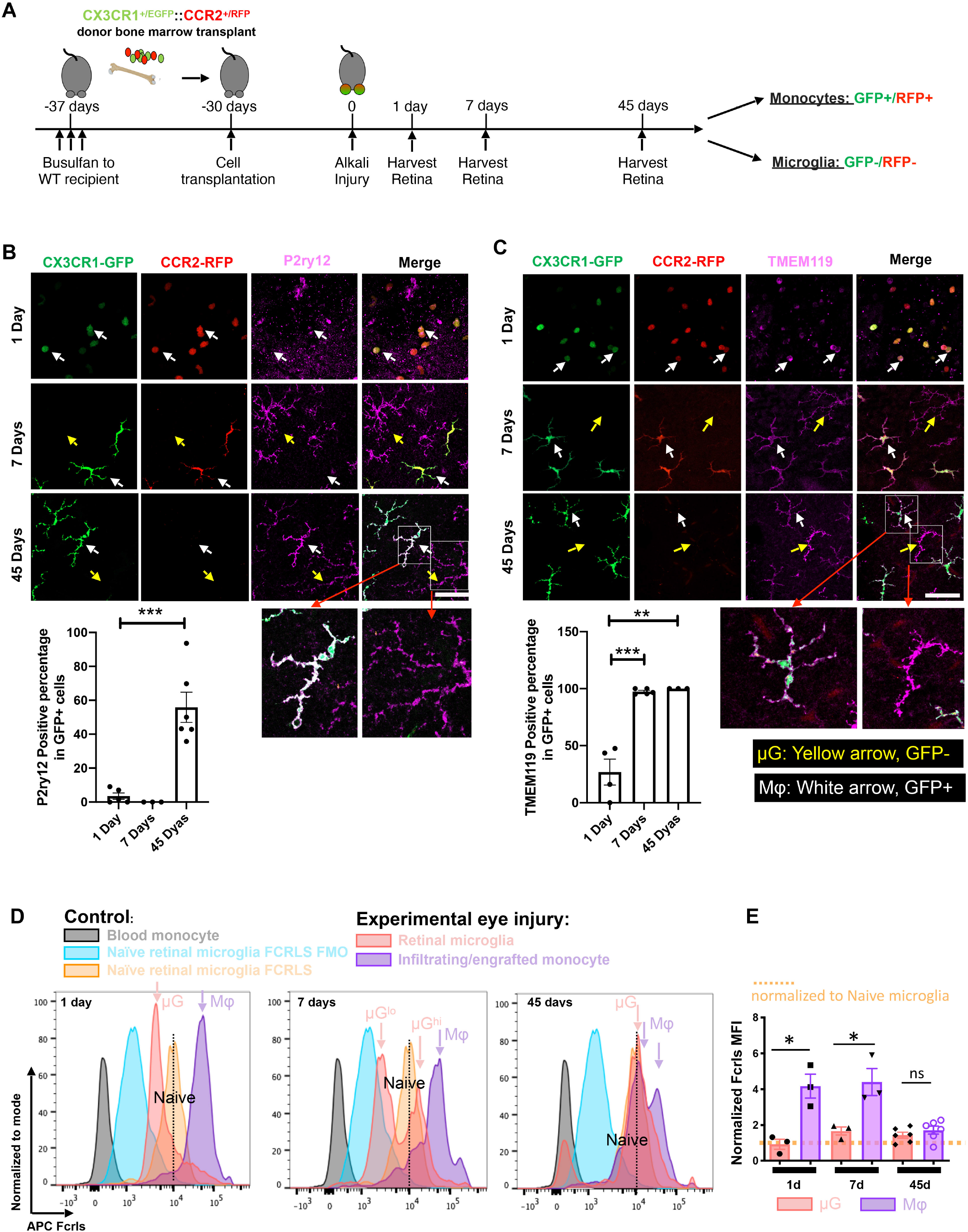
Acquisition of P2RY12, TMEM119, and FCRLS by donor-derived retinal myeloid cells. **(A)** CX3CR1^+/EGFP^::CCR2^+/RFP^ bone marrow chimeras were used to distinguish EGFP^+^ donor-derived cells from EGFP−host cells. CCR2−RFP expression varies after engraftment and is not required to remain positive in donor-derived cells. **(B)** P2RY12 was rare or undetectable in donor-derived retinal cells at day 1, undetectable at day 7, and detected in approximately 55% at day 45. **(C)** TMEM119 was detected in approximately 25% of donor-derived cells at day 1 and approximately 100% at days 7 and 45. White arrows indicate donor-derived cells; yellow arrows indicate host microglia. Scale bars, 50 µm. **(D, E)** FCRLS fluorescence distributions and MFI were measured separately by flow cytometry. Anti-CX3CR1–PE-Cy7 distinguished host EGFP− from donor EGFP^+^ cells; FCRLS was detected in the APC channel (see Fig. S6). FCRLS was low or undetectable in blood monocytes and increased in donor-derived retinal cells after injury, with elevated early fluorescence and lower intensity by day 45. Data are mean ± SEM; each dot represents one animal, with at least three animals per group. Reported comparisons used independent t-tests: *P < 0.05, **P < 0.01, ***P < 0.001. MFI, median fluorescence intensity; Mφ, macrophages; µG, microglia.

P2RY12 was rare or undetectable in donor-derived retinal cells at day 1 and undetectable at day 7, with expression evident by day 14 and present in approximately 55% of engrafted cells by day 45 **(Fig. 2B; Fig. S4)**. Donor-derived cells acquired a ramified morphology. P2RY12 expression persisted at 16 months in the CX3CR1^CreER-EYFP^::ROSA26tdTomato chimera model, with heterogeneity among animals **(Fig. S5A, B)**. TMEM119 was detected in approximately 25% of donor-derived cells at day 1 and in approximately 100% by day 7, with expression maintained at day 45 and 16 months **(Fig. 2C; Fig. S5C)**. Host EGFP^−^ cells also expressed P2RY12 and TMEM119 during the study period **(Fig. 2B, C)**.

FCRLS was assessed by flow cytometry in CX3CR1^+/EGFP^::CCR2^+/RFP^ chimeras. Anti-CX3CR1–PE-Cy7 distinguished CX3CR1^+^ host EGFP^−^ cells from donor EGFP^+^ cells, with the gating strategy shown in Figure S6. A fluorescence-minus-one control defined the FCRLS fluorescence background. FCRLS fluorescence was higher in donor-derived retinal cells than in blood monocytes by day 1 and remained detectable at day 45 **(Fig. 2D, E)**. The pan-myeloid marker IBA1 was detected in approximately 85% of donor-derived cells at day 1 and approximately 100% at days 7 and 45; host microglia also expressed IBA1 **(Fig. S7)**.

We also examined donor-derived retinal cells after NaIO3 injury, which damages the retinal pigment epithelium [17]. The illustrative day 14 images showed acquisition of P2RY12, TMEM119, and IBA1. At three months, TMEM119 and IBA1 were detected in approximately 100% of EGFP^+^ cells, P2RY12 in approximately 60%, and MHC-II in approximately 50% **(Fig. S8)**. Thus, marker acquisition was also observed in this injury model, although P2RY12 expression remained heterogeneous.

Donor-derived cells were also detected in the brain and spinal cord of busulfan-conditioned chimeras that received no additional CNS injury. At one month, these cells expressed TMEM119 and IBA1, whereas P2RY12 expression varied by CNS region. At one year, P2RY12 was detectable in both regions **(Fig. S9, S10)**. The cells had ramified morphology and no detectable MHC-II under the staining conditions used.

### Chromatin accessibility changes accompany acquisition of microglia-associated features

To examine chromatin changes associated with engraftment [18–20], we performed ATAC-seq on sorted circulating monocytes, naive microglia, injured host microglia, and engrafted monocytes. Chromatin accessibility differed among these populations **(Fig. 3)**. At *P2ry12* and *Tmem119*, regions accessible in microglia but not in circulating monocytes became accessible in engrafted cells **(Fig. 3B, D)**. Regions at *Fcrls* and *Aif1* were already accessible in circulating monocytes **(Fig. 3C, E)**. These findings indicate locus-specific chromatin remodeling accompanying retinal engraftment.

**Figure 3.**
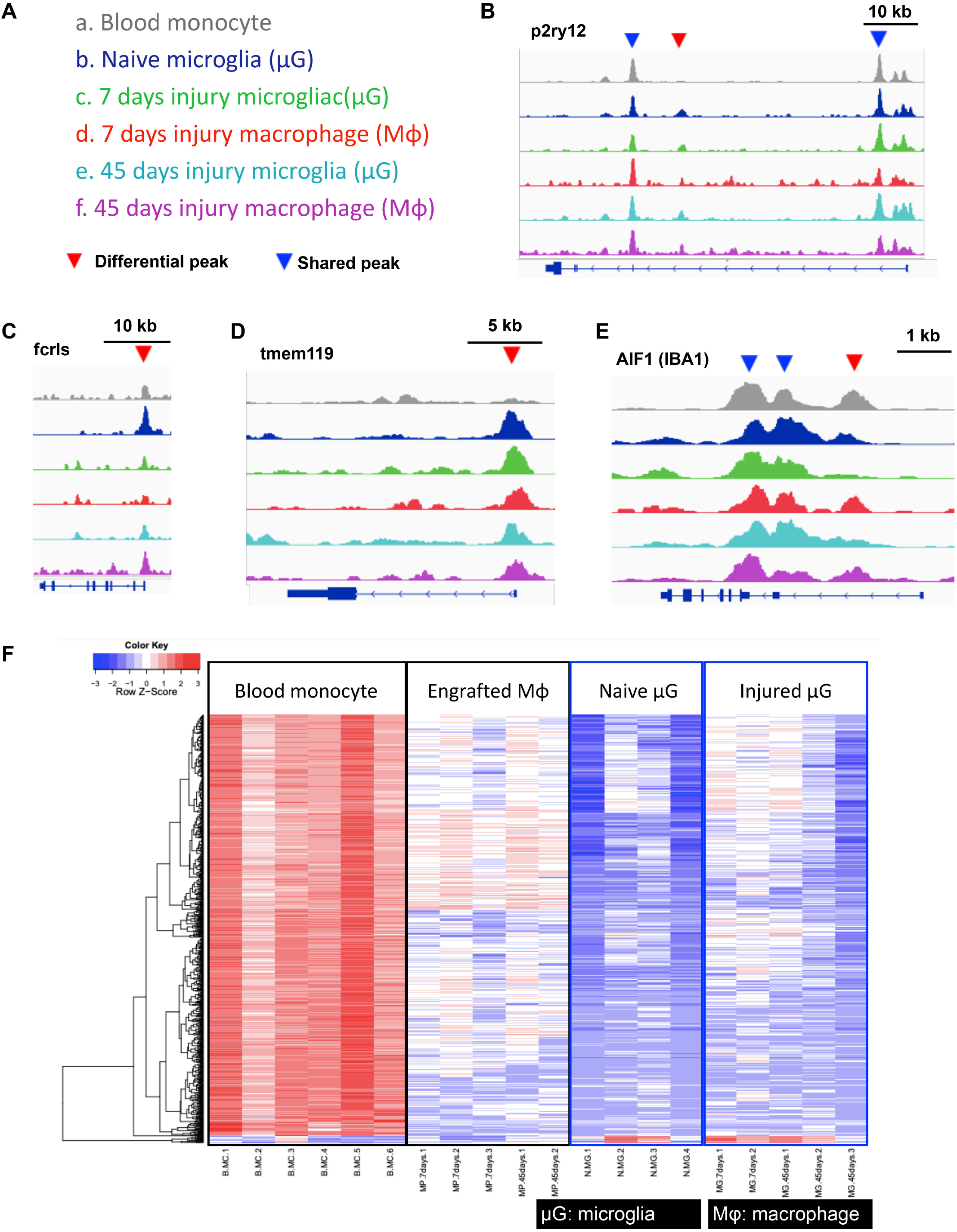
Chromatin accessibility in circulating monocytes and retinal myeloid populations. **(A)** Color key for the ATAC-seq groups. **(B)** *P2ry12* accessibility tracks include a region present in microglia and acquired by donor-derived cells after retinal engraftment (red arrowhead); other accessible regions are shared with circulating monocytes (blue arrowheads). **(C)** *Fcrls* accessibility is observed across the sampled populations. **(D)** *Tmem119* includes a region accessible in microglia and engrafted cells but not in circulating monocytes. (E) *Aif1*, encoding IBA1, shows accessible regions across the groups. **(F)** Heatmap of differential chromatin-accessibility peaks, scaled by row as Z-scores. Rows represent genomic peaks and columns represent biological samples arranged by population. Row clustering is shown; no sample-clustering dendrogram is displayed. Circulating monocytes, n = 6 samples; engrafted cells, n = 5 (three at day 7 and two at day 45); naive microglia, n = 4; injured host microglia, n = 5 (two at day 7 and three at day 45). Each ATAC-seq sample pooled five retinas from five mice. The heatmap shows both shared and distinct accessibility patterns.

The heatmap of differential ATAC-seq peaks showed changes after engraftment as well as persistent differences among circulating monocytes, engrafted monocytes, and host microglia **(Fig. 3F)**. Motif analysis identified shared and time-dependent enrichment relative to circulating monocytes (Fig. 4). PU.1 and AP-1 motifs were enriched in naive microglia and in engrafted cells at both time points. CTCF and RUNX enrichment varied with time. MAF and MITF motifs were enriched in engrafted cells, whereas NFKB1 enrichment was detected at day 7. By contrast, CEBP, IRF2, and ATF3 motifs met the enrichment threshold in naive microglia but not in the engrafted populations. Together, these patterns identify candidate regulatory programs associated with retinal engraftment.

**Figure 4.**
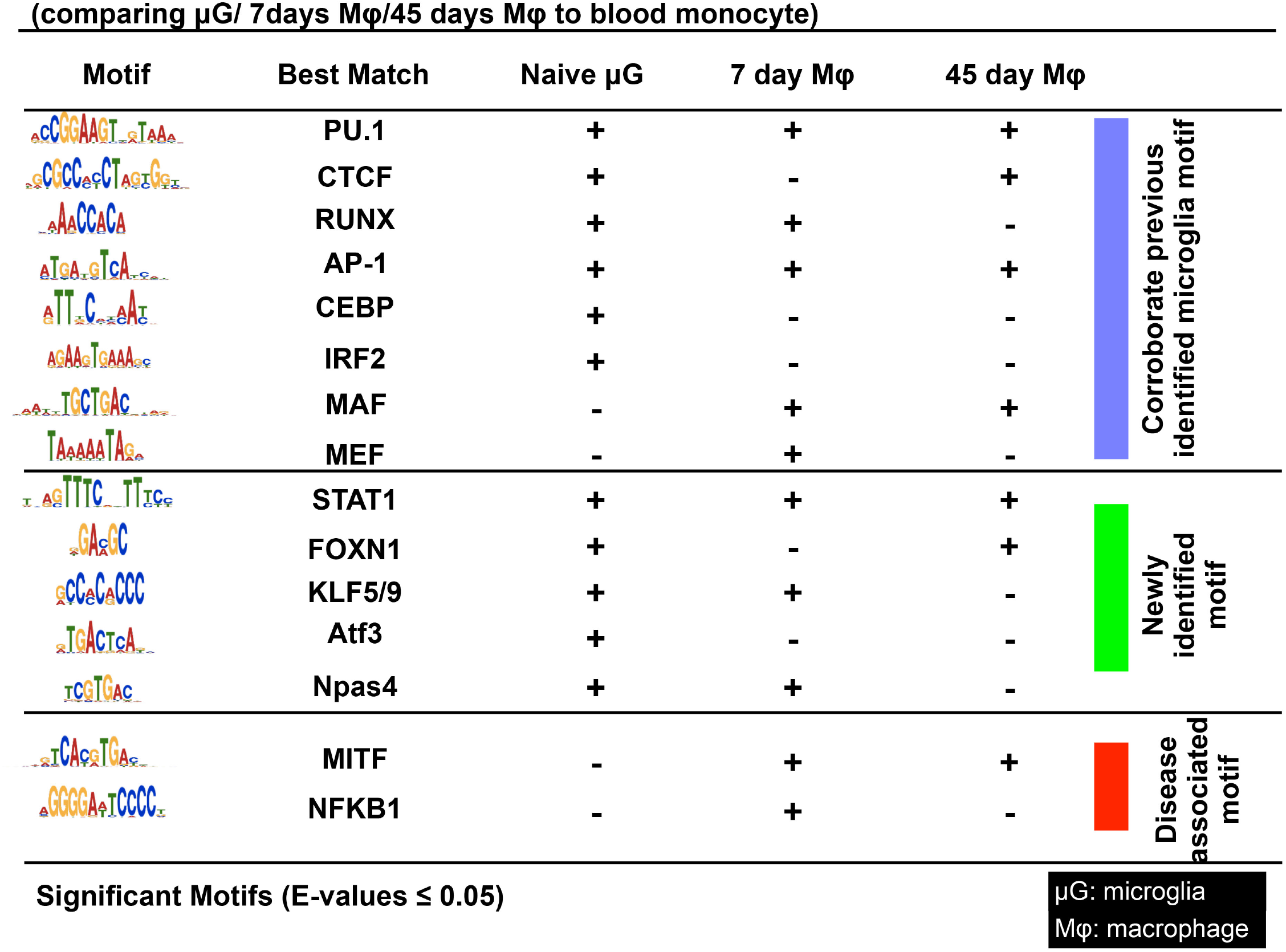
Transcription-factor motif enrichment in accessible chromatin. Motif enrichment in naive microglia and engrafted retinal myeloid cells at days 7 and 45 was evaluated relative to circulating monocytes. A plus sign denotes enrichment meeting the displayed E-value threshold (≤0.05); a minus sign denotes failure to meet that threshold. PU.1, AP-1, and STAT1 motifs were enriched in all three populations. CTCF and FOXN1 were enriched in naive microglia and day-45 engrafted cells; RUNX, KLF5/9, and Npas4 in naive microglia and day-7 engrafted cells. CEBP, IRF2, and ATF3 met the enrichment threshold only in naive microglia. MAF and MITF were enriched in engrafted cells at both times, whereas MEF and NFKB1 were enriched only at day 7. Colored sections organize the displayed motif groups.

### Engrafted retinal monocytes share transcriptional features with host microglia but retain differences at one year

We performed bulk RNA-seq on sorted naive microglia, engrafted monocytes, and host microglia from injured retinas, with three replicates per population at one year after injury (Fig. 5). Differentially expressed genes (DEGs) were defined by |log₂ fold change| > 2 and adjusted P < 0.05. Principal component analysis showed that PC1, PC2, and PC3 accounted for 39%, 23%, and 13% of the variance, respectively. One engrafted-monocyte replicate was separated from the other engrafted samples along PC1. Complementary PC1–PC2 and PC2–PC3 projections were examined to characterize the distribution of samples **(Fig. S11A)**. In the PC2–PC3 projection, naive microglia formed a distinct group, separated from injured host microglia and engrafted monocytes predominantly along PC3, while the two injury-associated populations showed greater between-replicate dispersion.

**Figure 5.**
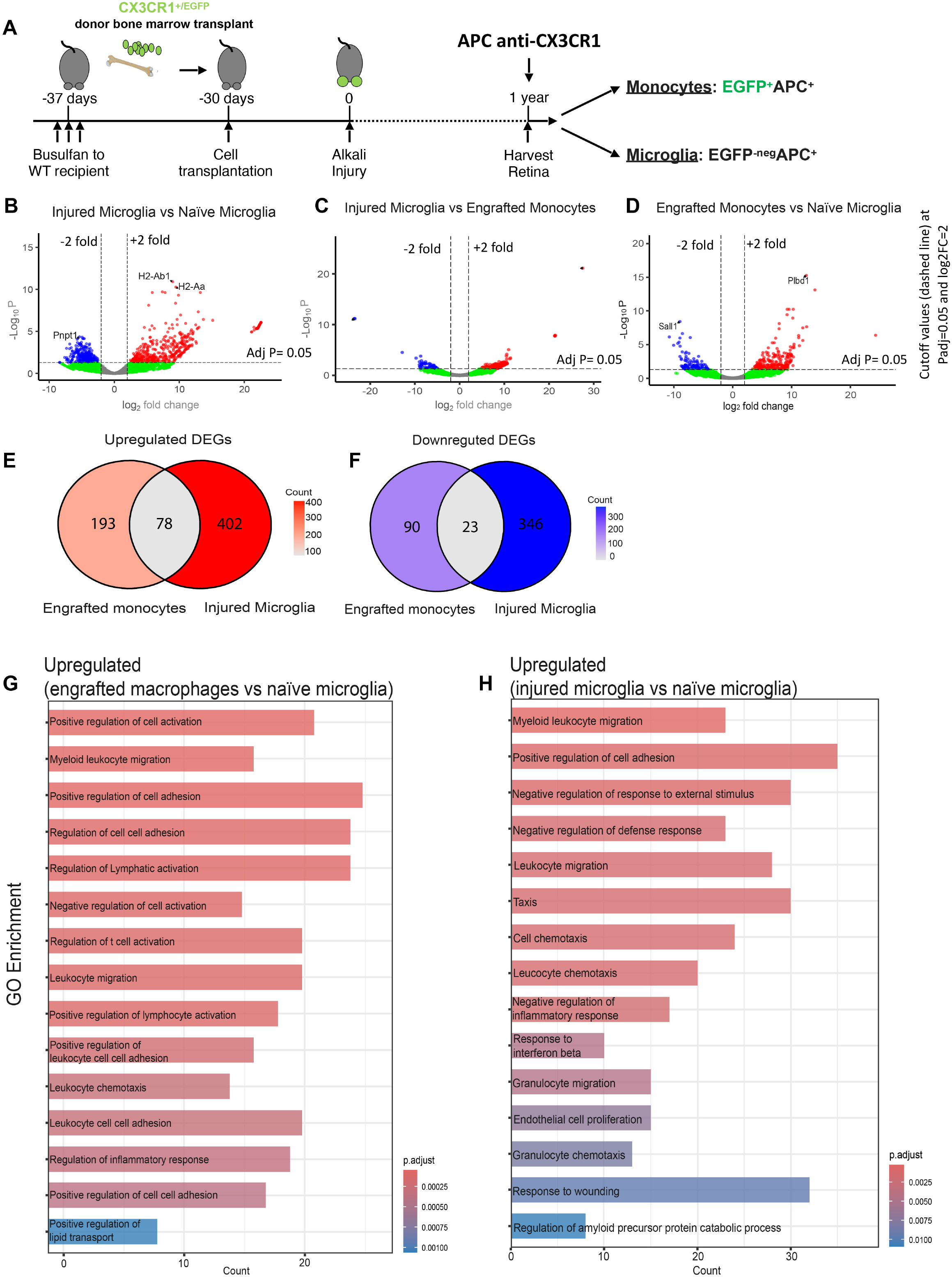
Bulk RNA-seq comparisons of retinal myeloid populations one year after injury. **(A)** Experimental timeline and donor-versus-host identification using CX3CR1^+/EGFP^ bone marrow chimeras and anti-CX3CR1–APC. **(B–D)** Volcano plots for injured versus naive microglia **(B)**, injured microglia versus engrafted monocytes **(C)**, and engrafted monocytes versus naive microglia **(D)**. Positive log2 fold change indicates higher expression in the first-named population. Significance criteria were |log2 fold change| > 2 and adjusted P < 0.05. Red and blue indicate significant increases and decreases, respectively; green indicates genes exceeding the fold-change threshold without meeting the significance threshold. Selected genes are annotated. **(E, F)** Overlap of upregulated **(E)** and downregulated **(F)** DEGs in injured microglia and engrafted monocytes, each relative to naive microglia. The comparisons share 78 upregulated and 23 downregulated genes. **(G, H)** The top 15 displayed Gene Ontology biological-process terms for genes upregulated in engrafted monocytes **(G)** or injured microglia **(H)**, each relative to naive microglia. Bar length indicates the number of associated DEGs and color denotes the adjusted P value.

Among 10937 genes in the injured-versus-naive microglia comparison, 480 were significantly upregulated and 369 downregulated. When injured microglia were compared with engrafted monocytes, 177 genes were significantly upregulated and 71 downregulated. The engrafted-monocyte-versus-naive microglia comparison included 10,800 genes, of which 271 were significantly upregulated and 113 downregulated **(Fig. 5B–D)**. These comparisons reveal shared and distinct transcriptional features in the two injury-associated populations.

Genes upregulated in injured microglia relative to naive microglia included the MHC-II genes *H2-Ab1*, *H2-Aa*, and *H2-Eb1*, together with *Cd74* **(Fig. 5B)**. Chemokine-associated genes (*Ccl8*, *Ccl7*, *Cxcl2*, *Cxcl10*, *Cxcl13*, and *Cxcl16*) and interferon-responsive genes (*Ifi47*, *Ifi44*, *Ifi203*, *Ifit3*, *Ifitm2*, and *Ifitm3*) were also upregulated. The most significantly downregulated gene, *Pnpt1* **(Fig. 5B)**, encodes a polyribonucleotide nucleotidyltransferase involved in mitochondrial RNA processing [21, 22] and it has been implicated in inflammasome regulation in macrophages [23].

In injured microglia compared with engrafted monocytes, *Gm14268* and *Pgm5* were the most significant up- and downregulated genes, respectively **(Fig. 5C)**. *Gm14268* is a long noncoding transcript reported in studies of photoreceptor differentiation [24], whereas *Pgm5* encodes phosphoglucomutase-like protein 5. *Tlr8*, *Ptpn22*, *Rnasel*, *Serpine1*, *Serpina3g*, *Itgal*, and *Itga4* also had higher expression in engrafted monocytes than in injured microglia. These expression differences identify candidate distinctions in immune signaling and tissue interactions.

*Plbd1* was the most significant upregulated gene in engrafted monocytes relative to naive microglia, whereas *Sall1*, a regulator of microglial identity [25, 26], was the most significantly downregulated gene **(Fig. 5D)**. Reduced *Sall1* expression persisted at one year despite acquisition of other microglia-associated markers. Several MHC-II- and chemokine-associated genes were upregulated in both engrafted monocytes and injured microglia relative to naive microglia. The two comparisons shared 78 upregulated DEGs and 23 downregulated DEGs **(Fig. 5E, F)**.

Gene Ontology analysis of genes upregulated relative to naive microglia identified overlapping but distinct sets of enriched biological processes **(Fig. 5G, H)**. In engrafted monocytes, prominent terms concerned cell activation, lymphocyte activation, adhesion, leukocyte migration, and regulation of inflammatory responses **(Fig. 5G).** In injured microglia, enriched terms included leukocyte migration and chemotaxis, response to interferon-β, negative regulation of inflammatory responses, and response to wounding **(Fig. 5H).**

We compared genes upregulated in either engrafted monocytes or injured microglia relative to naive microglia with a preselected reference list of 104 NF-κB-associated genes derived from the Gilmore laboratory repository (https://www.bu.edu/nf-kb/gene-resources/target-genes/). Eight distinct genes met the threshold criteria in at least one comparison: *Cd74*, *Il2ra*, *Ptx3*, *S100a6*, *Il1b*, *Mmp9*, *Ccr7*, and *Cd69* **(Fig. S11B, C)**. The heatmap summarizes row-wise standardized mean expression for each population.

### Engrafted monocytes downregulate Ly6C and CD45 in subsets

We next assessed the stability of conventional myeloid markers after retinal engraftment. Ly6C^+^ monocytes are recruited during CNS inflammation [7, 27] and after inducible microglial depletion [10]. CD45 intensity is also commonly used to distinguish resident microglia from infiltrating myeloid cells [28], although this distinction may vary with cell state and time.

In CX3CR1^+/EGFP^::CCR2^+/RFP^ chimeras, donor-derived retinal cells showed a progressive change in reporter phenotype over 45 days **(Fig. 6A, B)**. Early cells were predominantly CCR2^high^CX3CR1^low^ (Group 1), whereas the major population at day 45 was CCR2^low/−^CX3CR1^high^ (Group 5), overlapping the marker profile of host microglia (Group 6).

**Figure 6.**
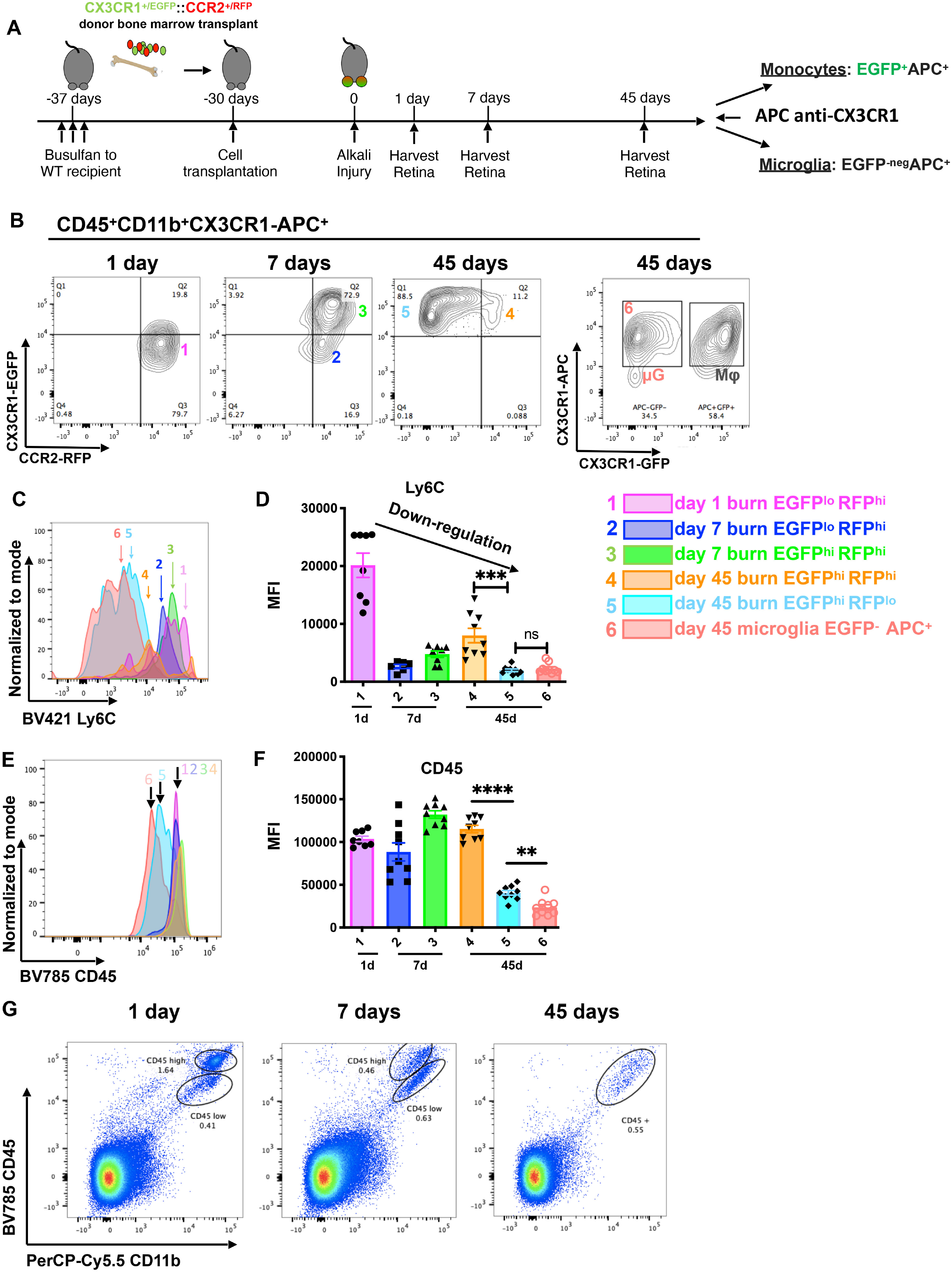
Ly6C and CD45 expression in host and donor-derived retinal myeloid cells. **(A)** CX3CR1^+/EGFP^::CCR2^+/RFP^ bone marrow chimera model and flow-cytometric identification of host cells (EGFP−CX3CR1-APC+) and donor-derived cells (EGFP+CX3CR1-APC+). **(B)** Reporter-defined donor populations (Groups 1–5) at days 1, 7, and 45, and host microglia (Group 6). The 88.5% label in the day-45 reporter plot is a representative quadrant value. **(C, D)** Ly6C fluorescence distributions and MFI. The major day-45 engrafted population has lower Ly6C than early populations, while a CCR2-high subset retains higher expression. The displayed Group 5 versus Group 6 comparison is nonsignificant. **(E, F)** CD45 fluorescence distributions and MFI. CD45 decreases in the major engrafted population but remains significantly different from host microglia in the displayed Group 5 versus Group 6 comparison. **(G)** Representative CD45/CD11b plots show increasing overlap of donor-cell and host-cell intensity ranges. Each dot represents one animal; at least eight animals per group. Comparisons are reported as independent t-tests. ns, not significant; **P < 0.01, ***P < 0.001, ****P < 0.0001. MFI, median fluorescence intensity.

Ly6C expression was high in early CCR2^high^ donor-derived cells and lower in the major CCR2^low/−^CX3CR1^high^ population at day 45 **(Group 5; Fig. 6C, D)**. Lund and colleagues likewise reported reduced Ly6C expression after CNS engraftment in a microglial depletion model [10]. A smaller CCR2^high^ population retained high Ly6C at day 45 (Group 4), demonstrating persistent phenotypic heterogeneity **(Fig. 6C, D)**.

CD45 expression also varied over time. CD45^high^ donor-derived cells remained evident at days 1 and 7. By day 45, many donor-derived cells shifted toward intermediate CD45 expression **(Fig. 6G)**. However, CD45 intensity remained significantly different between the major engrafted population and host microglia (Groups 5 and 6; **Fig. 6F**), and a higher-expressing donor subset persisted **(Fig. 6E–G)**.

Thus, Ly6C and CD45 intensities overlap between subsets of long-term engrafted cells and host microglia in this model. These markers alone cannot reliably assign origin to every retinal myeloid cell after prolonged engraftment. Figure 7 summarizes the observed changes in morphology and marker expression.

**Figure 7.**
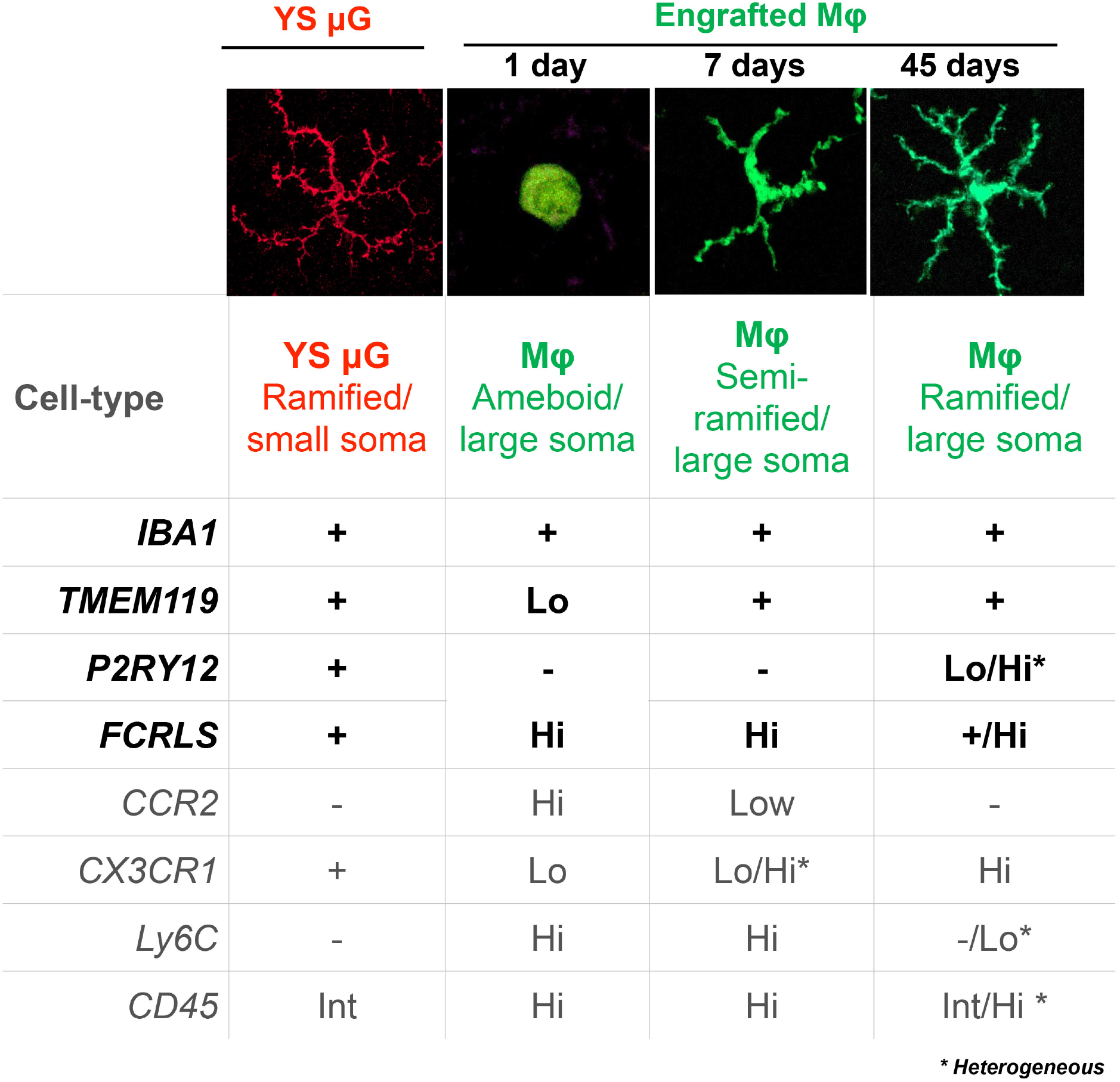
Summary of morphology and marker changes during retinal engraftment. Donor-derived retinal cells become more ramified over time, with reduced CCR2 and Ly6C and lower CD45 in subsets, together with acquisition of TMEM119, P2RY12, and FCRLS. IBA1 is a pan-myeloid marker. P2RY12 and other features remain heterogeneous, as indicated. The schematic integrates the morphological and marker-expression findings from the separate assays and summarizes predominant patterns; individual cells and subsets vary. Percentages indicate marker-positive fractions. YS µG, yolk-sac-derived microglia; Mφ, macrophages.

## Discussion

This study advances our understanding of retinal monocyte engraftment by defining the temporal acquisition of microglial markers and the accompanying changes in chromatin accessibility and transcription. Our previous work established that infiltrating monocytes persist in the retina and remain proinflammatory despite adopting a microglia-like morphology [2]. Here, we show that this resemblance extends to acquisition of P2RY12, TMEM119, and FCRLS with distinct kinetics, together with reduced Ly6C and CD45 expression in subsets. These phenotypic changes are accompanied by molecular adaptation that includes both shared features and persistent differences relative to host microglia. By integrating temporal, phenotypic, and molecular analyses, our findings extend previous descriptions of microglia-like adaptation by monocyte-derived CNS macrophages [2, 8–12] and clarify how engrafted cells become increasingly difficult to distinguish from resident microglia.

The timing of marker acquisition helps explain this identification challenge. TMEM119 was detected soon after retinal entry, whereas P2RY12 appeared later and remained heterogeneous. FCRLS fluorescence also increased in donor-derived retinal cells. Adaptation continued over longer periods in other CNS compartments: in the spinal cord of conditioned chimeras without additional CNS injury, P2RY12 was undetectable at one month but detectable at one year. Thus, the ability of individual markers to distinguish myeloid populations changes with tissue, time, and experimental context [6, 7, 11]. P2RY12 and TMEM119 reflect aspects of cell state as well as identity, while IBA1 is broadly expressed by myeloid cells; their expression also varies across developmental, inflammatory, and neurodegenerative settings [29–37]. These observations support interpreting marker profiles alongside cellular origin and anatomical location, particularly when assessing the contribution of long-term engrafted cells to retinal inflammation.

Chromatin-accessibility profiling revealed regulatory changes accompanying this phenotypic adaptation. Increased accessibility at *P2ry12* and *Tmem119* in engrafted cells was consistent with acquisition of the corresponding markers and with evidence that the tissue environment shapes myeloid gene regulation [9, 38, 39]. The broader motif-enrichment patterns showed that adaptation involved both common and population-associated regulatory features. Shared PU.1 and AP-1 enrichment suggested overlapping regulatory programs, whereas differences in MAF, MITF, and NFKB1 motif enrichment identified candidate programs associated with engraftment. The chromatin data therefore extend the phenotypic observations by showing that acquisition of microglial features occurs within a regulatory landscape that remains distinguishable from that of host microglia.

The one-year bulk RNA-seq analysis provided a complementary view of the long-term state of these populations. Both engrafted monocytes and injured host microglia upregulated genes associated with MHC-II antigen presentation and immune signaling relative to naive microglia. Their direct comparison nevertheless identified differences in immune-signaling and adhesion-associated transcripts. GO enrichment highlighted cell activation, adhesion, leukocyte migration, and inflammatory-response programs, while analysis of NF-κB-associated genes identified shared upregulation of *Cd74* and *Il2ra* alongside differences in other genes. These profiles show that long-term retinal residence and acquisition of microglial markers coexist with sustained inflammatory gene expression. They also indicate that the response to injury includes shared transcriptional features without eliminating differences between engrafted cells and host microglia.

The persistence of these differences is consistent with earlier studies showing that peripheral myeloid cells can occupy a depleted CNS macrophage niche while retaining transcriptomic, epigenomic, or functional properties distinct from those of resident microglia [9–12]. *Sall1* provides an informative example. Kim and colleagues reported *Sall1* expression and extensive transcriptional adaptation in monocyte-derived cells in the aging brain [40]. Their transplantation experiments also detected *Sall1* transcripts, although engrafted brain populations remained transcriptionally distinguishable from yolk-sac-derived microglia at 32 weeks. In our injured-retina model, *Sall1* expression remained significantly lower in engrafted cells than in naive microglia at one year. Given the role of SALL1 in regulating microglial identity [25, 26], this reduction marks a substantive molecular distinction despite acquisition of P2RY12, TMEM119, and FCRLS. Differences in tissue, injury, and conditioning provide important context for comparing the extent of adaptation across studies. This context is also relevant to microglial replacement, which has shown benefit in CSF1R-associated microgliopathy [41] and is supported by experimental strategies for replacing CNS myeloid populations [42]. Our findings contribute a retinal perspective by defining molecular features that persist after prolonged engraftment.

The conditions permitting engraftment are equally important when relating these findings to microglial maintenance. Ajami and colleagues showed that local self-renewal sustains adult CNS microglia [43] and that infiltrating monocytes promote experimental autoimmune encephalomyelitis progression without contributing to the resident microglial pool during remission [44]. Mildner and colleagues further demonstrated that adult brain engraftment depends on defined host conditions [45], while parabiosis provides a complementary approach to assessing circulating-cell contributions without marrow conditioning [46]. Previous studies using this busulfan-conditioning approach found no widespread engraftment of donor-derived CX3CR1^+^ cells in the uninjured retinal parenchyma, consistent with preservation of the blood–retinal barrier [2, 3]. In our busulfan-conditioned bone marrow chimeras, donor-derived myeloid cells engrafted in the brain and spinal cord without additional CNS injury, whereas retinal engraftment was examined after injury. The brain and spinal cord observations therefore extend the finding of marker acquisition to additional CNS tissues within the transplantation model, while underscoring the importance of anatomical and experimental setting.

Several features of this model define the scope of the molecular comparisons. The reporter strategy identifies donor origin rather than a permanent, mature-monocyte-specific lineage, with “engrafted monocytes” denoting the donor-derived retinal population studied here. Retinal border-associated macrophages were not separately excluded during sorting or resolved by a dedicated classification of the scRNA-seq data [47], leaving their contribution to the molecular profiles unresolved. Chromatin and expression measurements sampled different stages of adaptation and provide complementary views of this process. The identified regulatory programs offer candidates for functional investigation, while targeted chromatin methods such as CUT&RUN could further characterize histone modifications and transcription factor occupancy [48].

Together, these findings show that retinal engraftment involves coordinated phenotypic and molecular adaptation while preserving differences from host microglia. The resulting overlap in commonly used markers makes cellular-origin information particularly important for identifying the populations involved in persistent retinal inflammation and interpreting their responses to therapeutic intervention.

## Materials and Methods

### Animal Models and Procedures

All animal experiments followed the Association for Research in Vision and Ophthalmology Statement for the Use of Animals in Ophthalmic and Vision Research and were approved by the Animal Care Committee of Massachusetts Eye and Ear. Mice were bred and housed at the Massachusetts Eye and Ear Animal Facility and entered the experiments at 6–12 weeks of age.

### Bone-Marrow Chimera Generation

Bone marrow transplant chimeras were generated to distinguish donor-derived from host myeloid cells. Recipient C57BL/6J mice received intraperitoneal busulfan (35 mg/kg; Sigma-Aldrich) 7, 5, and 3 days before transplantation. Donor bone marrow cells (5 × 10⁶), obtained from CX3CR1^+/EGFP^::CCR2^+/RFP^, CX3CR1^+/EGFP^, or CX3CR1^CreER-EYFP^::ROSA26tdTomato mice, were administered by tail vein injection one month before experimental injury. Mice received trimethoprim/sulfamethoxazole (Bactrim) in drinking water for 15 days after busulfan treatment. Recipient C57BL/6J (stock 000664), *Ccr2*-RFP (017586), CX3CR1-EGFP (005582), *Cx3cr1*-CreER (021160), and ROSA26-tdTomato (007914) mice were obtained from The Jackson Laboratory. CX3CR1^+/EGFP^::CCR2^+/RFP^ donor mice were generated by crossing the *Ccr2*-RFP [49] and CX3CR1-EGFP lines.

### Tamoxifen administration

In the CX3CR1^CreER-EYFP^::ROSA26tdTomato chimera model, donor-derived CX3CR1-expressing cells were labeled by administering tamoxifen two weeks after corneal injury. Tamoxifen (20 mg/mL in corn oil) was given by two intraperitoneal injections of 75 mg/kg separated by one day, as described previously [50].

### Mouse Model of Alkali Burn

One month after bone marrow transplantation, corneal alkali burns were induced as described previously [13]. Mice were anesthetized with ketamine (60 mg/kg) and xylazine (6 mg/kg). After topical proparacaine was applied, a 2-mm filter paper soaked in 1 M NaOH was placed on the cornea for 20 seconds. The eye was immediately irrigated with sterile saline for 15 minutes. Postprocedural treatment included subcutaneous Ethiqa XR (buprenorphine, 3.25 mg/kg; Covetrus North America, FP-001) and topical Polytrim (polymyxin B/trimethoprim; Bausch & Lomb).

### NaIO3 injury model

Retinal pigment epithelial injury was induced by a single intraperitoneal injection of NaIO3 (30 mg/kg; Sigma-Aldrich, S4007). Tissues were analyzed 14 days or three months after injection, as indicated in Figure S8.

### Flow Cytometry and Cell Sorting

Flow cytometry was used to measure FCRLS, Ly6C, and CD45 and to isolate retinal myeloid populations. Retinas were dissociated with papain (Worthington Biochemical Corporation, LK003150). In reporter bone marrow chimeras, donor-derived cells were identified as CD45^+^CD11b^+^EGFP^+^ and host cells as CD45^+^CD11b^+^CX3CR1^+^EGFP^−^. The anti-CX3CR1 fluorophore varied by assay, as shown in the corresponding figure. After Fc-receptor blocking with anti-CD16/32 (clone 2.4G2), cells were stained with the antibodies listed in Table 1, analyzed or sorted on a BD FACSAria III, and evaluated with FlowJo 10.

**Table 1.** Antibodies used for flow cytometry and immunostaining. Both P2RY12 antibodies were used in the experimental investigations. Dashes indicate details not specified in the reagent information.

| ANTIBODY | FLUOROPHORE/SECONDARY | BRAND | CLONE | CATALOG<br>NUMBER | DILUTION | APPLICATION |
| --- | --- | --- | --- | --- | --- | --- |
| CD45 | Brilliant Violet 785 | Biolegend | 30-F11 | 103149 | 1 to 100 | Flow cytometry |
| CD11B | PerCP-Cyanine 5.5 | Biolegend | M1/70 | 101228 | 1 to 100 | Flow cytometry |
| LY6C | Brilliant Violet 421 | Biolegend | HK1.4 | 128031 | 1 to 100 | Flow cytometry |
| CX3CR1 | APC | Biolegend | SA011F11 | 149008 | 1 to 100 | Flow cytometry |
| CX3CR1 | Brilliant Violet 605 | Biolegend | SA011F11 | 149027 | 1 to 100 | Flow cytometry |
| FCRLS | Alexa Fluor 647 donkey anti rat | Dr. Margeta |  |  | 1 to 200 | Flow cytometry |
| IBA1 | Alexa Fluor 647 donkey anti rabbit | Wako |  | 019-19741 | 1 to 300 | Immunostaining |
| TMEM119 | Alexa Fluor 647 donkey anti guinea pig | Synaptic systems |  | 400 004 | 1 to 300 | Immunostaining |
| P2RY12 | Alexa Fluor 647 donkey anti rabbit | Dr. Margeta |  |  | 1 to 200 | Immunostaining |
| P2RY12 | Alexa Fluor 647 donkey anti rabbit | Anaspec |  | as-55043a | 1 to 200 | Immunostaining |
| MHC-II | Alexa Fluor 647 donkey anti rat | BD Bioscience | M5/114.15.2 | 556999 | 1 to 100 | Immunostaining |

### Single-cell RNA sequencing and gene expression profiling

Flow-sorted CD45^+^CD11b^+^ cells were encapsulated in microdroplets for single-cell library preparation. Barcoded hydrogel beads and reverse-transcription reagents were introduced using a high-throughput microfluidic platform to initiate in-droplet cDNA synthesis. Libraries were prepared according to Klein et al. [51] and sequenced on an Illumina HiSeq 2500 with 100-bp paired-end reads and approximately 1,000 cells per sample. Reads were mapped to the Mus musculus GRCm38/mm10 reference genome, quantified as gene counts, and normalized to reads per million.

### RNA isolation and quantitative real-time PCR

A bone marrow transplantation model distinguished donor-derived retinal cells from host microglia. Naive retinal microglia and blood monocytes were collected from uninjured chimeras; host microglia and engrafted monocytes were collected 45 days after ocular injury. Sorted cells were collected directly into 1 mL TRIzol (Thermo Fisher Scientific, 15596026), and RNA was extracted according to the manufacturer’s instructions. GlycoBlue Coprecipitant (Thermo Fisher Scientific, AM9515) was added during RNA precipitation. Reverse transcription and cDNA amplification used the SMART-Seq v4 Ultra Low Input RNA Kit (Takara, 634890). RNA from approximately 400 cells yielded approximately 30 ng cDNA after 18 amplification cycles. cDNA was quantified with the Qubit dsDNA High Sensitivity Assay (Thermo Fisher Scientific, Q32851). Quantitative PCR used TaqMan probes, TaqMan Universal PCR Master Mix (Thermo Fisher Scientific, 4304437), and 250–500 pg cDNA per reaction.

### ATAC-seq

The CX3CR1^+/EGFP^::CCR2^+/RFP^ bone marrow chimera model was used to isolate donor-derived CD45^+^CD11b^+^EGFP^+^ cells and host CD45^+^CD11b^+^CX3CR1^+^EGFP^−^ cells. Each sample pooled five retinas from five mice. Approximately 1,000–5,000 cells were processed using the Active Motif ATAC-seq kit (13150). Nuclei were isolated in 100 µL ice-cold ATAC lysis buffer and incubated with tagmentation master mix at 37°C and 800 rpm for 30 minutes. DNA was purified, and libraries were amplified for 13 PCR cycles using indexed primers. Library-fragment size distributions were assessed before sequencing. Sequencing on an Illumina HiSeq 2000 generated approximately 30 million 50-bp paired-end reads per sample. Reads were mapped to the mm9 mouse reference genome using BWA [52]. Fragments longer than 100 bp with both ends unambiguously mapped were retained. Hotspot2 (https://github.com/Altius/hotspot2) identified peaks at a false-discovery-rate threshold of 0.05 [53]. Peak overlap required at least 50% reciprocal overlap. Differential accessibility between replicate groups was evaluated using DiffBind, and motif enrichment was analyzed using MEME-ChIP.

### RNA-seq

Bulk RNA-seq compared naive microglia, engrafted monocytes, and host microglia from injured retinas at one year after corneal alkali injury, with three samples per population. **Figure 5A** depicts CX3CR1^+/EGFP^ donor bone marrow transplanted into conditioned C57BL/6J recipients, with EGFP and anti-CX3CR1–APC used to distinguish donor-derived and host cells. Noninjured eyes were used for the population designated naive microglia. Differential expression was analyzed with DESeq2 (1.50.2) after expression filtering, using |log2 fold change| > 2 and adjusted P-value< 0.05 as significance criteria. Volcano plots and Venn diagrams were generated with EnhancedVolcano (1.28.2) and VennDiagram (1.8.2), respectively. Enrichment analyses used clusterProfiler (4.18.4), with mouse gene sets obtained through msigdbr (26.1.0). The NF-κB-associated heatmap was generated with ComplexHeatmap (2.26.1) using mean expression across the three samples per population, standardized within each gene **(Fig. S11B)**.

### Immunohistochemistry and Imaging

Retinas were collected for flat-mount staining at the indicated time points after injury. Eyes were fixed overnight in 4% paraformaldehyde at 4°C and dissected. Brain and spinal cord tissues were fixed under the same conditions and sectioned at 12 µm. Tissues were blocked for one hour at room temperature in PBS containing 5% normal donkey serum and 0.25% Triton X-100, incubated with primary antibodies overnight or over a weekend at 4°C, and then incubated with secondary antibodies (Table 1). Retinas were flattened by radial incisions and mounted on glass slides with coverslips. Images were acquired with an Axio Imager M2 microscope (Zeiss, Oberkochen, Germany).

### Statistics

Statistical analyses were performed with GraphPad Prism 9. Median fluorescence intensity (MFI) was used for flow-cytometric quantification. Normality was assessed using the Shapiro–Wilk test, and group comparisons were evaluated using the tests indicated in the figure legends. P < 0.05 was considered statistically significant. RNA-seq differential-expression and enrichment analyses are described separately above.

## Acknowledgments

This work was supported by the Boston Keratoprosthesis Research Fund; the Department of Defense (W81XWH2210774 and W81XWH2010916); the National Institutes of Health (7R01EY013124 and 5P30EY003790); and the Retina Research Foundation.

---

**Supplementary Figure S1. Blood chimerism after busulfan conditioning and bone marrow transplantation**

**(A)** Timeline for assessing blood chimerism 30 days after transplantation. **(B)** Approximately 95% of blood CD45^+^CD11b^+^CX3CR1^+^ cells were donor reporter-positive across four mice. **(C)** Representative gating strategy distinguishing host from donor CX3CR1+ blood cells; 95.7% is the value in the illustrated donor-cell gate.

**Supplementary Figure S2. Selected transcripts in blood monocytes and retinal myeloid populations**

A CX3CR1^+/EGFP^ bone marrow chimera model distinguished donor-derived EGFP^+^ cells from host EGFP− cells within the CD45^+^CD11b^+^CX3CR1^−^BV605^+^ gate. Uninjured chimeras provided naive retinal microglia and blood monocytes; injured host microglia and engrafted cells were collected 45 days after corneal alkali injury. *P2ry12*, *Tmem119*, *Fcrls*, and *Aif1* transcripts were measured by quantitative PCR. *P2ry12*, *Tmem119*, and *Aif1* were normalized to blood monocytes; *Fcrls* was normalized to naive retinal microglia. Relative expression is displayed on logarithmic axes. Each dot represents one animal; n ≥ 3 per analysis.

**Supplementary Figure S3. Marker staining in circulating myeloid cells**

Circulating cells from CX3CR1^+/EGFP^::CCR2^+/RFP^ mice were stained for P2RY12, IBA1, TMEM119, and MHC-II. P2RY12, IBA1, and TMEM119 were not detected under the staining conditions shown, whereas MHC-II staining was evident. Cells varied in their CX3CR1-EGFP and CCR2-RFP reporter expression. Scale bars, 20 µm.

**Supplementary Figure S4. P2RY12 expression at 14 days after retinal engraftment**

Retinal flat mounts from CX3CR1^+/EGFP^::CCR2^+/RFP^ bone marrow chimeras show P2RY12 expression in donor-derived EGFP^+^ cells at day 14. The P2RY12-positive donor-derived cells illustrated have low or undetectable CCR2-RFP. Host microglia are donor-reporter-negative. Scale bar, 50 µm.

**Supplementary Figure S5. Marker expression 16 months after retinal engraftment**

**(A)** Wild-type recipients were conditioned with busulfan and transplanted with CX3CR1^CreER-EYFP^::ROSA26tdTomato donor marrow. Corneal injury was followed two weeks later by tamoxifen to label donor-derived CX3CR1-expressing cells. **(B–D)** P2RY12 (B), TMEM119 **(C)**, and IBA1 **(D)** staining at 16 months. P2RY12 positivity remained variable among animals; TMEM119 and IBA1 were detected in approximately all labeled cells in the displayed samples. White arrows indicate EYFP^+^tdTomato^+^ donor-derived cells; yellow arrows indicate reporter-negative host cells. n = 3 animals. Scale bars, 20 µm. BMT, bone marrow transplantation; TAM, tamoxifen; w, weeks; m, months.

**Supplementary Figure S6. Flow-cytometric gating for the FCRLS assay**

**(A)** Experimental timeline in bone marrow chimeras. **(B)** Retinal cells were selected using forward and side scatter, singlet gating, and exclusion of DAPI-positive cells. Within CD11b^+^CD45^+^ cells, anti-CX3CR1–PE-Cy7 identified CX3CR1^+^ host EGFP− and donor-derived EGFP^+^ populations.

**Supplementary Figure S7. IBA1 expression after retinal engraftment**

CX3CR1^+/EGFP^::CCR2^+/RFP^ bone marrow chimeras distinguish donor-derived EGFP^+^ retinal cells from host EGFP− cells. IBA1 was detected in approximately 85% of donor-derived cells at day 1 and approximately 100% at days 7 and 45. White arrows indicate donor-derived cells; yellow arrows indicate host cells. Each dot represents one animal, with at least three animals per group. ns, not significant by the reported independent t-test. Scale bars, 50 µm.

**Supplementary Figure S8. Marker expression after NaIO3-induced retinal injury**

**(A)** CX3CR1^+/EGFP^ bone marrow chimera model. White arrows indicate EGFP^+^ donor-derived cells; host cells are EGFP^−^. **(B)** Illustrative TMEM119, IBA1, P2RY12, and MHC-II staining 14 days after NaIO3 injury (n = 1 animal; scale bars, 50 µm). **(C)** Staining at three months (n = 4 animals; scale bars, 20 µm). Approximately 100% of donor-derived cells were TMEM119- or IBA1-positive, approximately 60% were P2RY12-positive, and approximately 50% were MHC-II-positive. Markers were assessed in the staining combinations shown.

**Supplementary Figure S9. Marker expression in brain and spinal cord one month after transplantation**

**(A)** CX3CR1^+/EGFP^ donor marrow was transplanted into busulfan-conditioned recipients. Brain and spinal cord were collected one month later without an additional CNS injury. **(B)** In brain, IBA1 and TMEM119 were detected in approximately all donor-derived EGFP^+^ cells, P2RY12 in approximately 50%, and MHC-II was not detected. **(C)** In spinal cord, IBA1 and TMEM119 were detected in approximately all donor-derived cells; P2RY12 and MHC-II were not detected. White arrows indicate donor-derived EGFP^+^ cells; yellow arrows indicate host EGFP− cells. n = 3 animals. Scale bars, 50 µm.

**Supplementary Figure S10. Marker expression in brain and spinal cord one year after transplantation**

**(A)** CX3CR1^+/EGFP^ donor marrow was transplanted into busulfan-conditioned recipients; brain and spinal cord were collected one year later without an additional CNS injury. **(B, C)** In brain **(B)** and spinal cord **(C)**, IBA1, TMEM119, and P2RY12 were detected in approximately all donor-derived EGFP^+^ cells; MHC-II was not detected under the staining conditions used. White arrows indicate donor-derived EGFP^+^ cells; yellow arrows indicate host EGFP^−^ cells. n = 3 animals. Scale bars, 50 µm.

**Supplementary Figure S11. Principal component analysis and NF-κB-associated transcription one year after monocyte engraftment**

**(A)** PCA of bulk RNA-seq samples from naive microglia, injured host microglia, and engrafted monocytes, with three samples per population. Complementary projections show PC1 versus PC2 (left) and PC2 versus PC3 (right). PC1, PC2, and PC3 explain 39%, 23%, and 13% of the variance, respectively. Each point represents one sample; connecting lines delineate samples within each population. Red, injured host microglia; green, engrafted monocytes; blue, naive microglia. **(B)** Heatmap of eight distinct NF-κB-associated genes upregulated relative to naive microglia in at least one population, using adjusted P < 0.05 and log₂ fold change > 2 as DEG criteria. Values are row-wise Z-scores of mean expression across three samples per population. Rows represent genes and columns represent population means; both were hierarchically clustered. **(C)** Corresponding DEG tables showing four upregulated genes in injured host microglia (*S100a6*, *Cd74*, *Il2ra*, and *Ptx3*) and six in engrafted monocytes (*Mmp9*, *Cd74*, *Il2ra*, *Il1b*, *Cd69*, and *Ccr7*), each compared with naive microglia. *Cd74* and *Il2ra* are shared between the two comparisons. Tables report DESeq2 baseMean, log2FoldChange, and adjusted P values.

## Data Availability Statement

Summary results are provided in the figures and supplementary materials.

## Author Contributions

J.L. and E.I.P. designed research; J.L., F.L., N.C. and B.Y. performed research; J.L., F.L., N.C., J.S., C.R., V.C., S.N., D.V., D.W., R.M. and M.A.M. analyzed data; and J.L., B.Y., C.R., J.C., D.E.M., D.V., D.W., R.M. and E.I.P. wrote the paper.

## Competing Interest Statement

The authors declare no competing interests.

## Notes

### Competing Interest Statement

The authors have declared no competing interest.

### Summary of Updates

This version adds bulk RNA sequencing of naive microglia, injured host microglia, and engrafted monocytes at one year after retinal injury. New analyses characterize shared and distinct transcriptional programs, including antigen presentation, inflammatory signaling, and reduced Sall1 expression in engrafted cells. Figure 5 and Supplementary Figure S11 present these results, including principal component analysis, differential gene expression, and functional enrichment analyses. The title, abstract, introduction, and discussion have been revised to integrate these findings and clarify the relationship between microglial marker acquisition, chromatin accessibility, and cellular origin. Methods and figure legends have been expanded, the transplantation and conditioning context clarified, and figures renumbered. The reference list, author list, and author contributions have also been updated.

