## Supplementary material for "Chromatin and transcriptional adaptation accompany monocyte engraftment in the injured retina": Sup Figures

### Sup. Figure 1

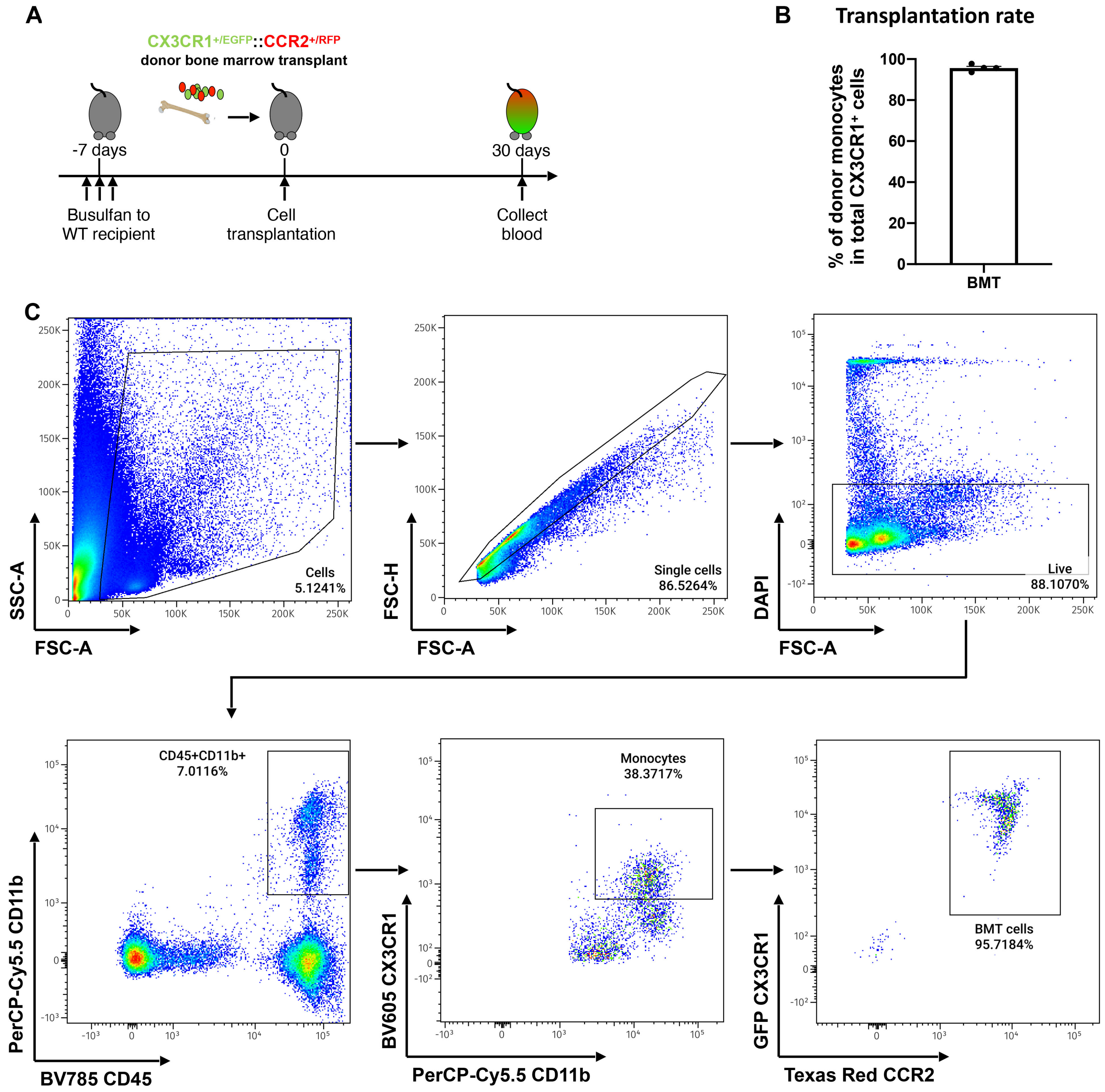

A
CX3CR1+/EGFP::CCR2+/RFP
donor bone marrow transplant
-7 days
0
30 days
Busulfan to WT recipient
Cell transplantation
Collect blood
B
Transplantation rate
% of donor monocytes in total CX3CR1+ cells
BMT
C
Cells 5.1241%
SSC-A
FSC-A
Single cells 86.5264%
FSC-H
FSC-A
Live 88.1070%
DAPI
FSC-A
CD45+CD11b+ 7.0116%
PerCP-Cy5.5 CD11b
BV785 CD45
Monocytes 38.3717%
BV605 CX3CR1
PerCP-Cy5.5 CD11b
BMT cells 95.7184%
GFP CX3CR1
Texas Red CCR2

**Sup. Figure 2**

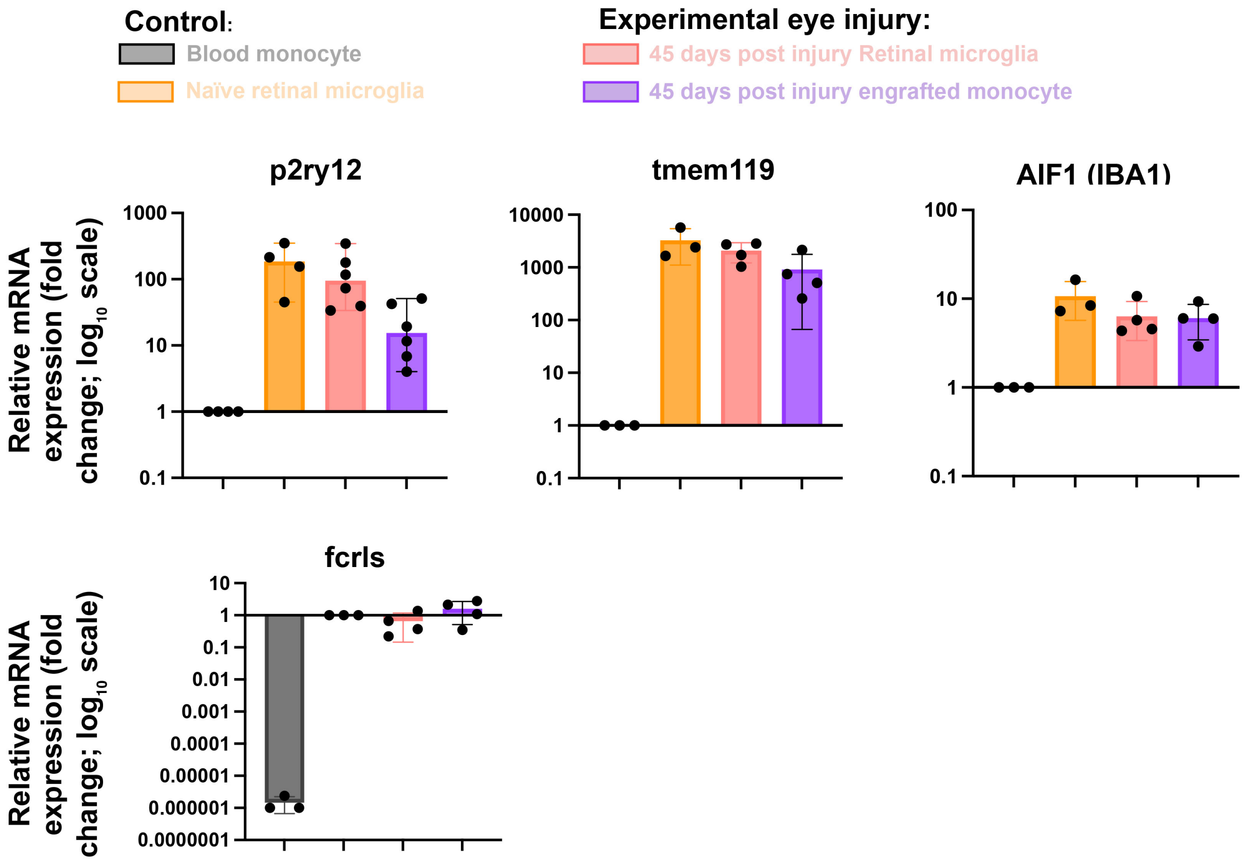
Control:
Blood monocyte
Naïve retinal microglia
Experimental eye injury:
45 days post injury Retinal microglia
45 days post injury engrafted monocyte
p2ry12
tmem119
AIF1 (IBA1)
fcrls
Relative mRNA expression (fold change; log10 scale)
1000
100
10
1
0.1
10000
0.01
0.001
0.0001
0.00001
0.000001
0.0000001

Sup. Figure 3

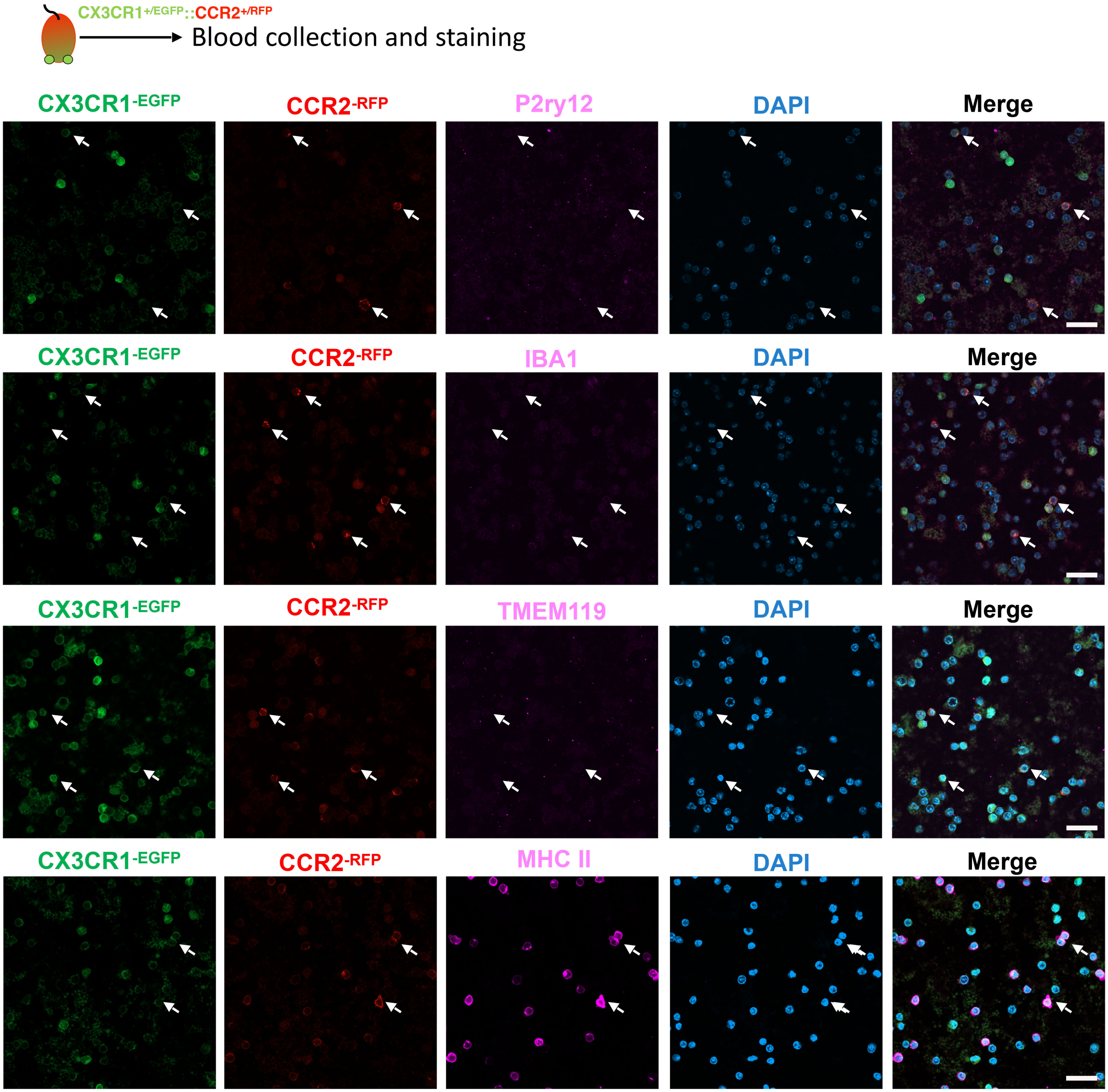

CX3CR1+/EGFP::CCR2+/RFP
Blood collection and staining
CX3CR1-EGFP
CCR2-RFP
P2ry12
DAPI
Merge
CX3CR1-EGFP
CCR2-RFP
IBA1
DAPI
Merge
CX3CR1-EGFP
CCR2-RFP
TMEM119
DAPI
Merge
CX3CR1-EGFP
CCR2-RFP
MHC II
DAPI
Merge

**Sup. Figure 4**

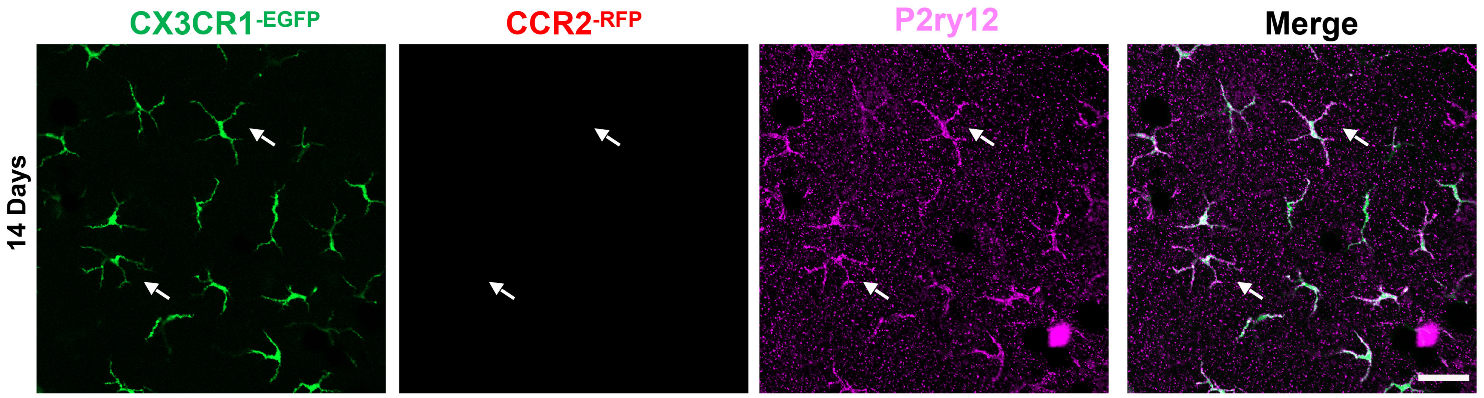

### Sup. Figure 5

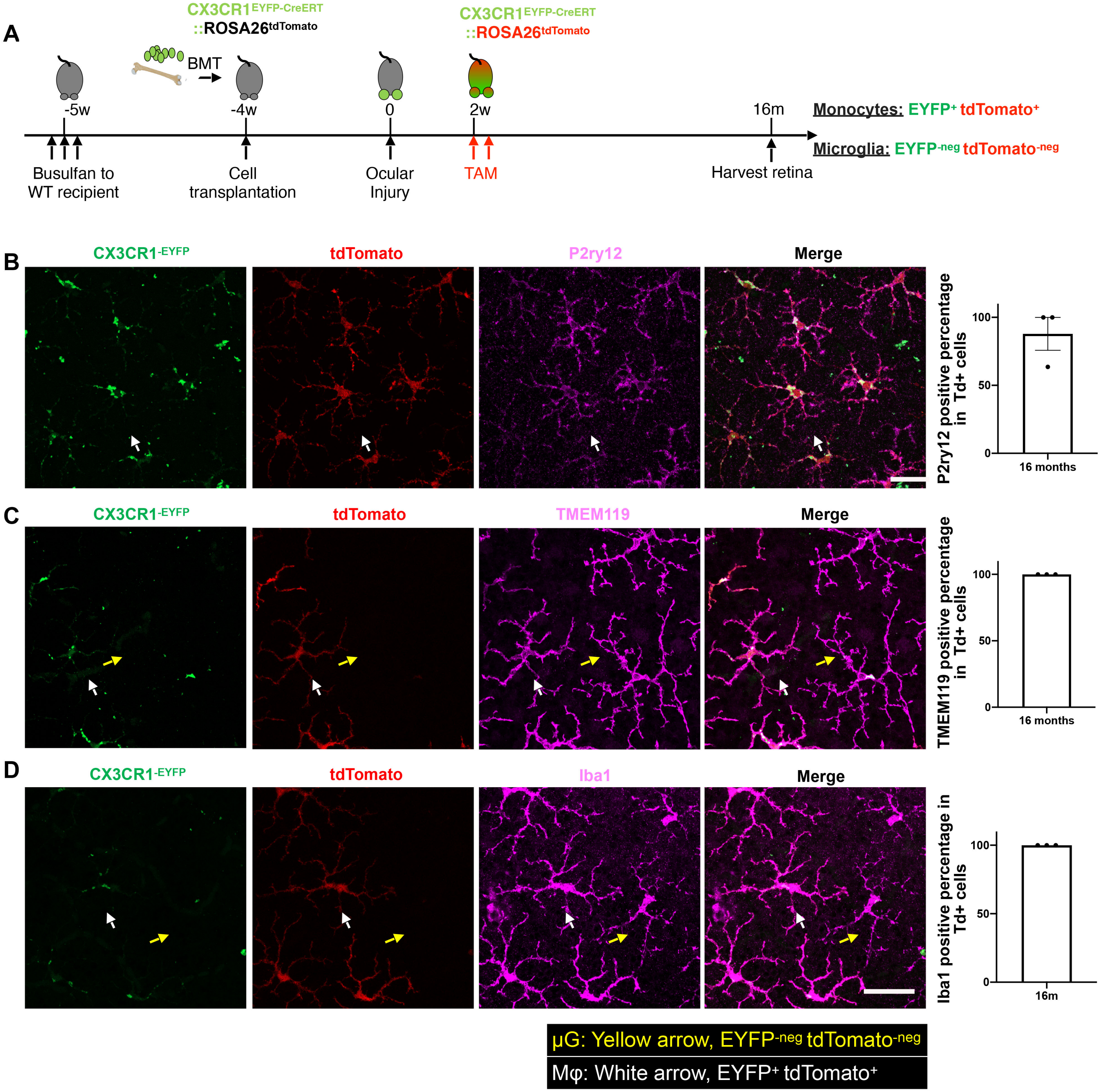
A
CX3CR1EYFP-CreERT ::ROSA26tdTomato
CX3CR1EYFP-CreERT ::ROSA26tdTomato
BMT
-5w
-4w
0
2w
16m
Busulfan to WT recipient
Cell transplantation
Ocular Injury
TAM
Harvest retina
Monocytes: EYFP+ tdTomato+
Microglia: EYFP-neg tdTomato-neg
B
CX3CR1-EYFP
tdTomato
P2ry12
Merge
P2ry12 positive percentage in Td+ cells
100
50
0
16 months
C
CX3CR1-EYFP
tdTomato
TMEM119
Merge
TMEM119 positive percentage in Td+ cells
100
50
0
16 months
D
CX3CR1-EYFP
tdTomato
Iba1
Merge
Iba1 positive percentage in Td+ cells
100
50
0
16m
μG: Yellow arrow, EYFP-neg tdTomato-neg
Mφ: White arrow, EYFP+ tdTomato+

### Sup. Figure 6

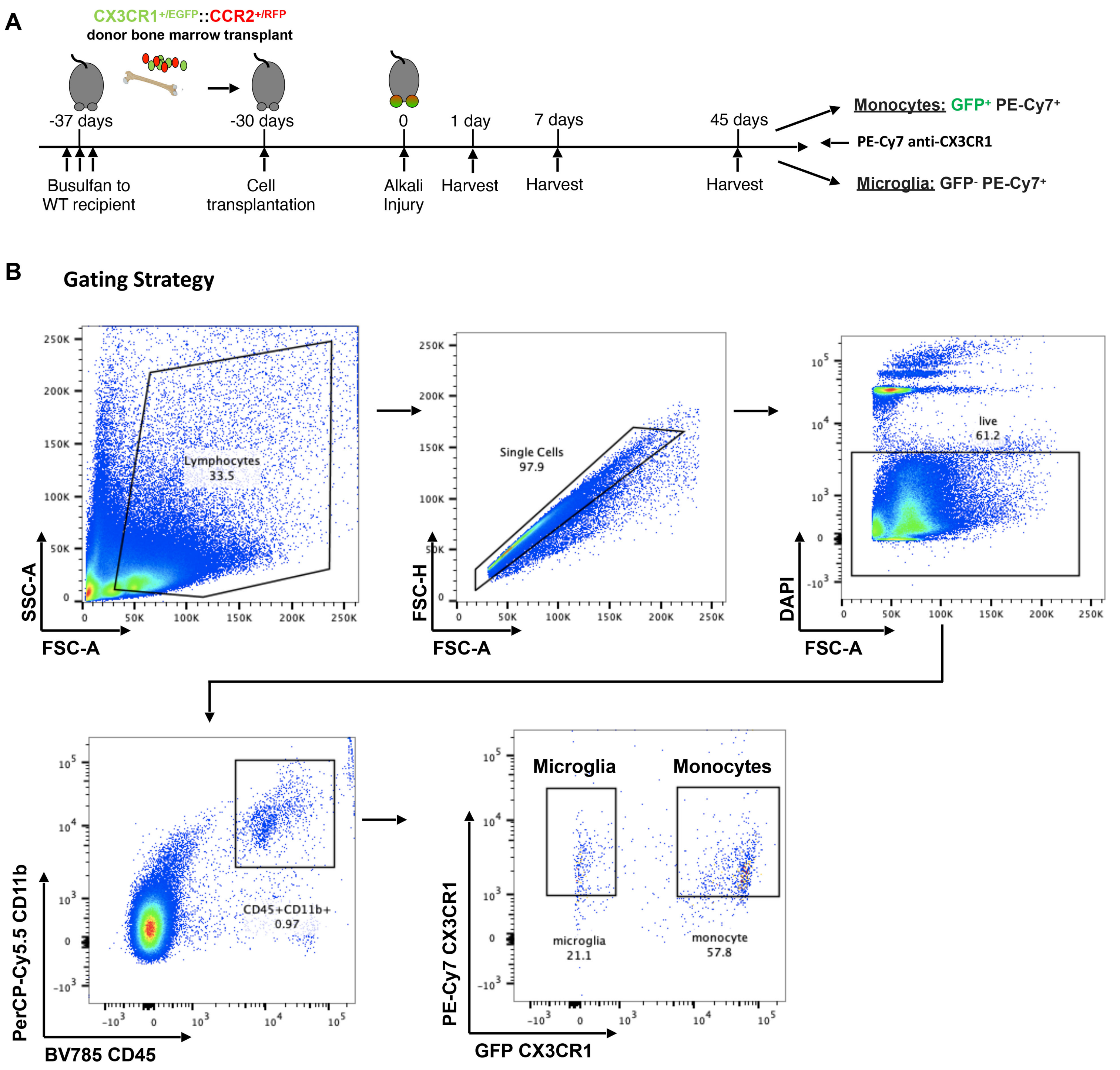
A
CX3CR1+/EGFP::CCR2+/RFP
donor bone marrow transplant
-37 days
-30 days
0
1 day
7 days
45 days
Busulfan to WT recipient
Cell transplantation
Alkali Injury
Harvest
Harvest
Harvest
Monocytes: GFP+ PE-Cy7+
PE-Cy7 anti-CX3CR1
Microglia: GFP- PE-Cy7+
B
Gating Strategy
Lymphocytes 33.5
SSC-A
FSC-A
Single Cells 97.9
FSC-H
FSC-A
live 61.2
DAPI
FSC-A
CD45+CD11b+ 0.97
PerCP-Cy5.5 CD11b
BV785 CD45
Microglia
Monocytes
microglia 21.1
monocyte 57.8
PE-Cy7 CX3CR1
GFP CX3CR1

### Sup. Figure 7

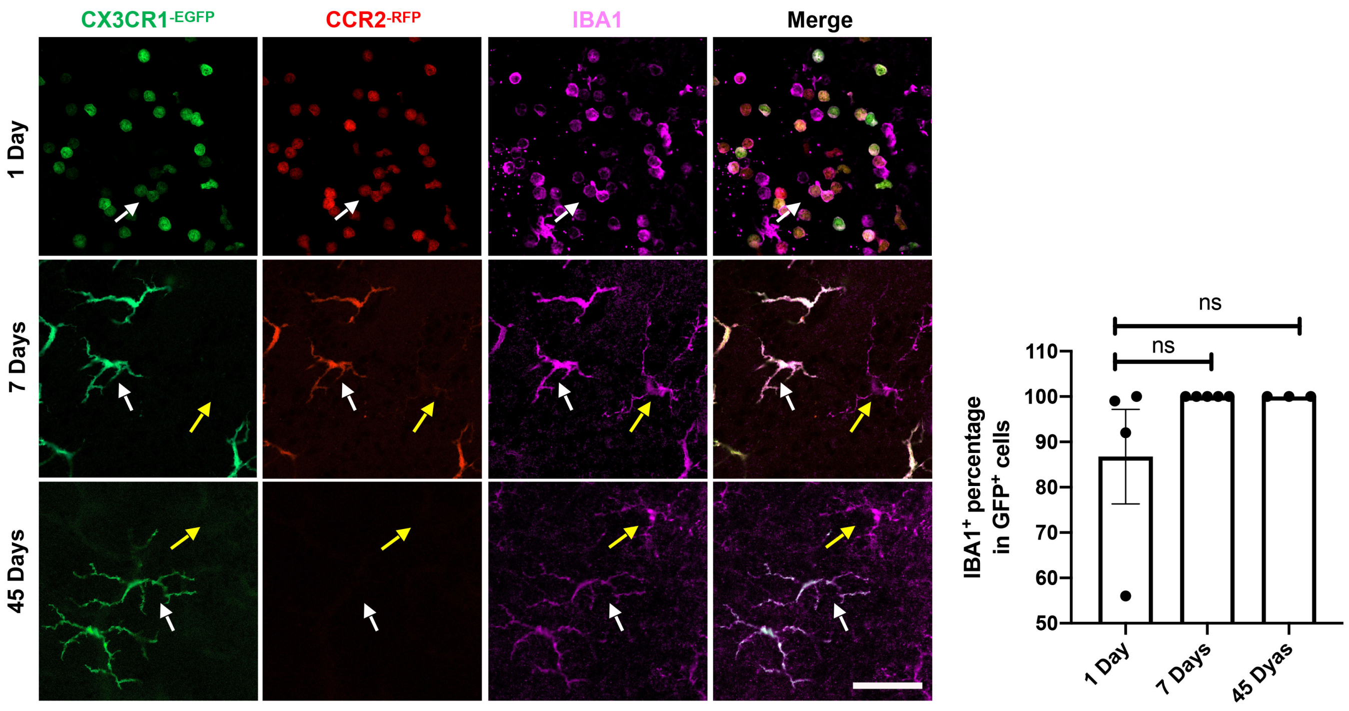
CX3CR1-EGFP
CCR2-RFP
IBA1
Merge
1 Day
7 Days
45 Days
ns
ns
110
100
90
80
70
60
50
IBA1+ percentage in GFP+ cells
1 Day
7 Days
45 Dyas

### Sup. Figure 8

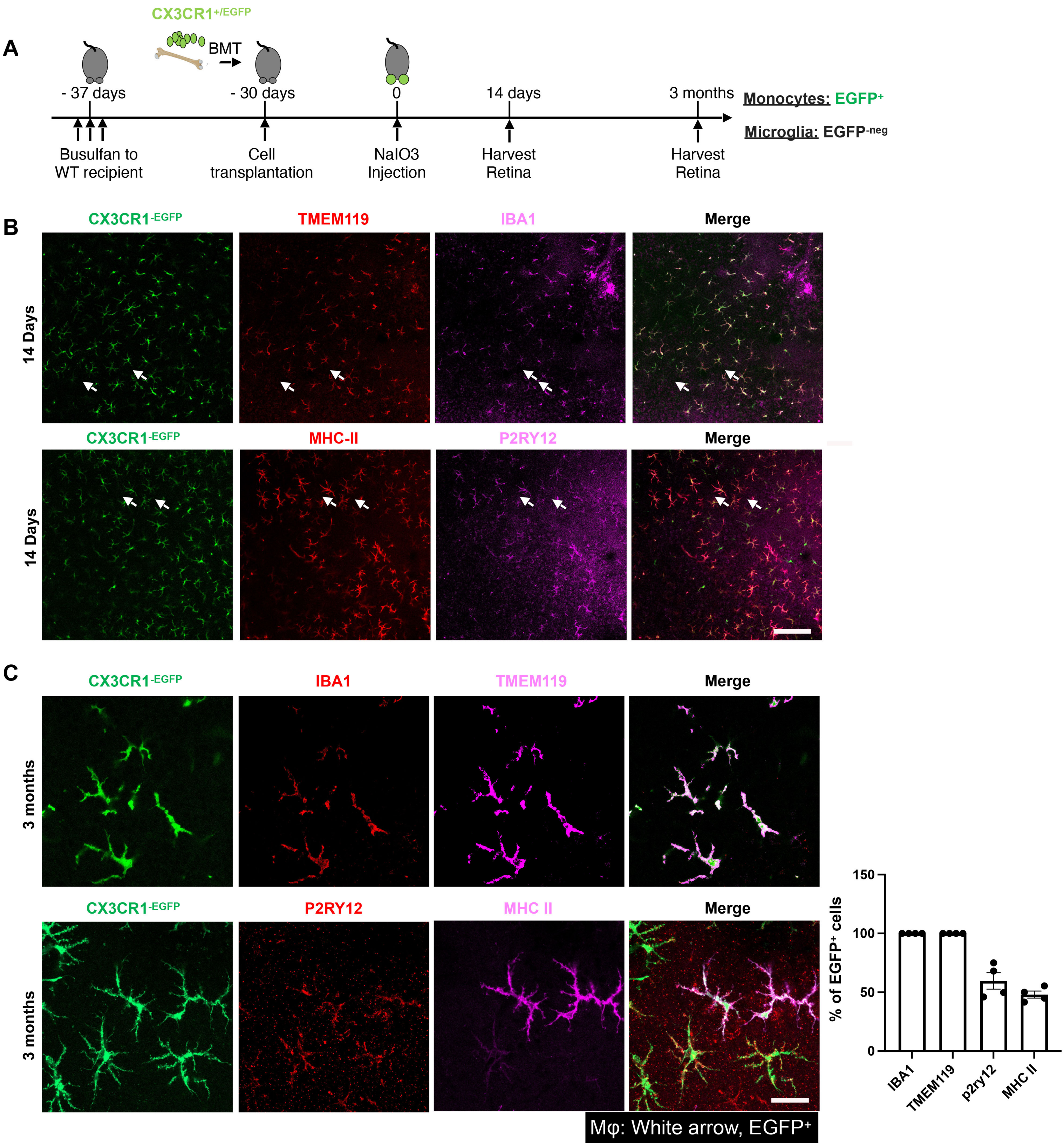
A
CX3CR1+/EGFP
BMT
- 37 days
- 30 days
0
14 days
3 months
Busulfan to WT recipient
Cell transplantation
NaIO3 Injection
Harvest Retina
Harvest Retina
Monocytes: EGFP+
Microglia: EGFP-neg
B
CX3CR1-EGFP
TMEM119
IBA1
Merge
14 Days
CX3CR1-EGFP
MHC-II
P2RY12
Merge
14 Days
C
CX3CR1-EGFP
IBA1
TMEM119
Merge
3 months
CX3CR1-EGFP
P2RY12
MHC II
Merge
3 months
Mφ: White arrow, EGFP+
% of EGFP+ cells
150
100
50
0
IBA1
TMEM119
p2ry12
MHC II

**Sup. Figure 9**

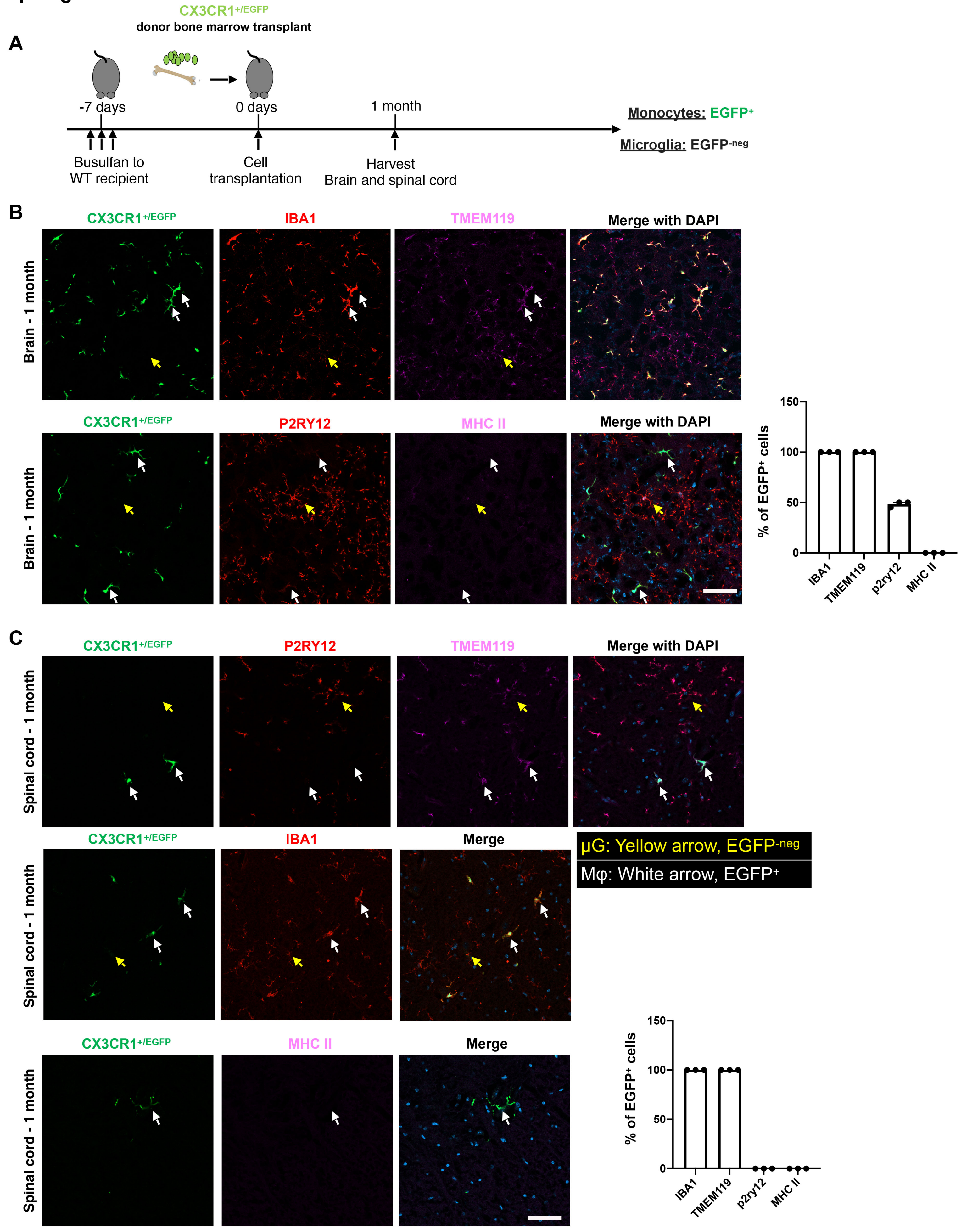
A
CX3CR1+/EGFP
donor bone marrow transplant
-7 days
0 days
1 month
Busulfan to WT recipient
Cell transplantation
Harvest Brain and spinal cord
Monocytes: EGFP+
Microglia: EGFP-neg
B
CX3CR1+/EGFP
IBA1
TMEM119
Merge with DAPI
Brain - 1 month
P2RY12
MHC II
% of EGFP+ cells
150
100
50
0
IBA1
TMEM119
p2ry12
MHC II
C
Spinal cord - 1 month
Merge
μG: Yellow arrow, EGFP-neg
Mφ: White arrow, EGFP+

### Sup. Figure 10

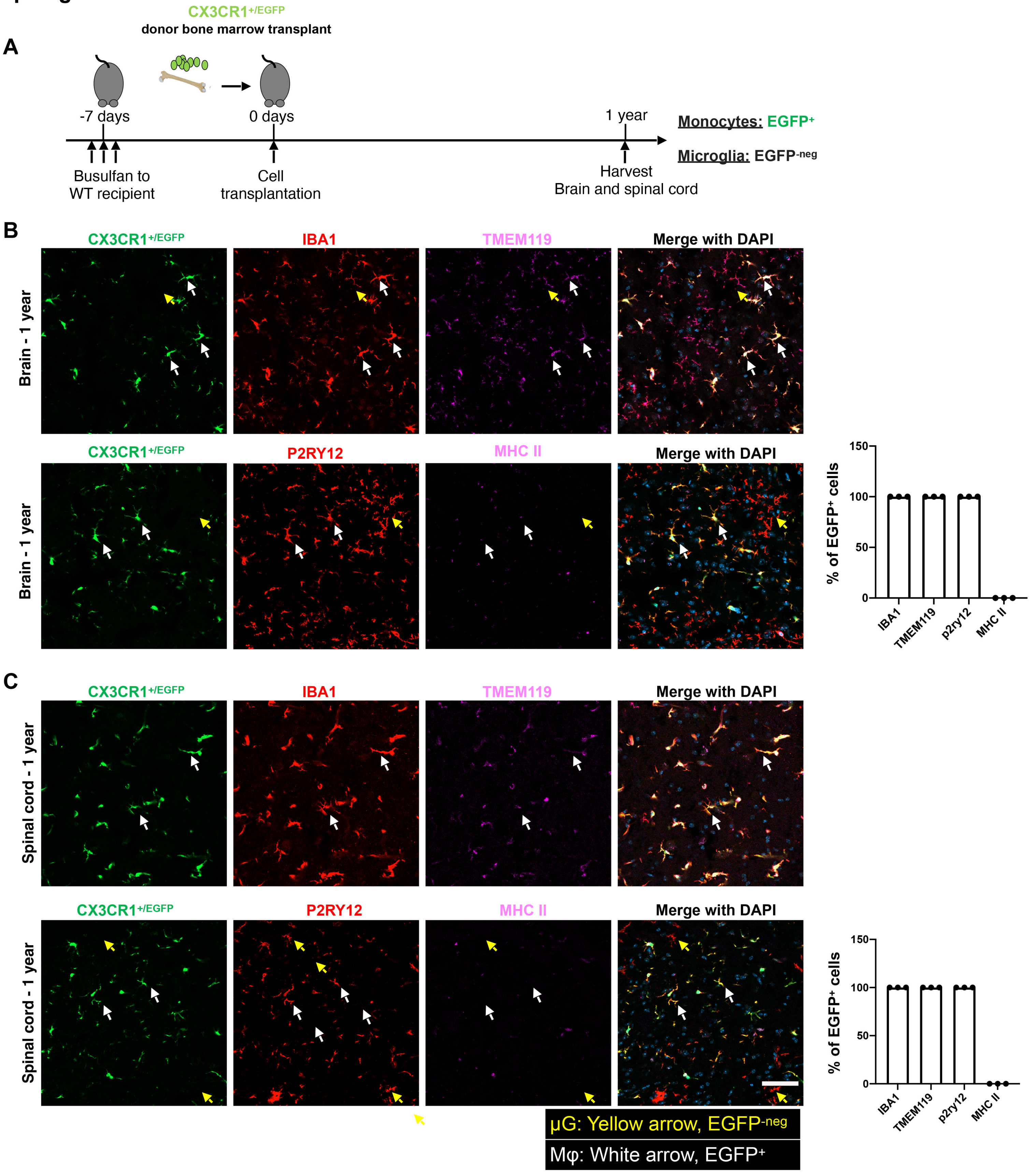

### Sup. Figure 11

Principal component analysis

**A**

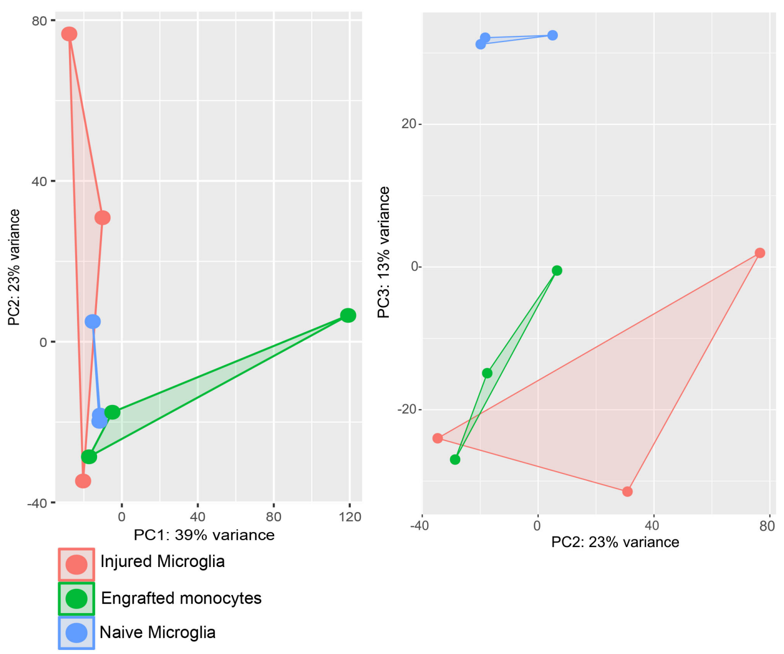


NF-κB-associated DEGs

**B**

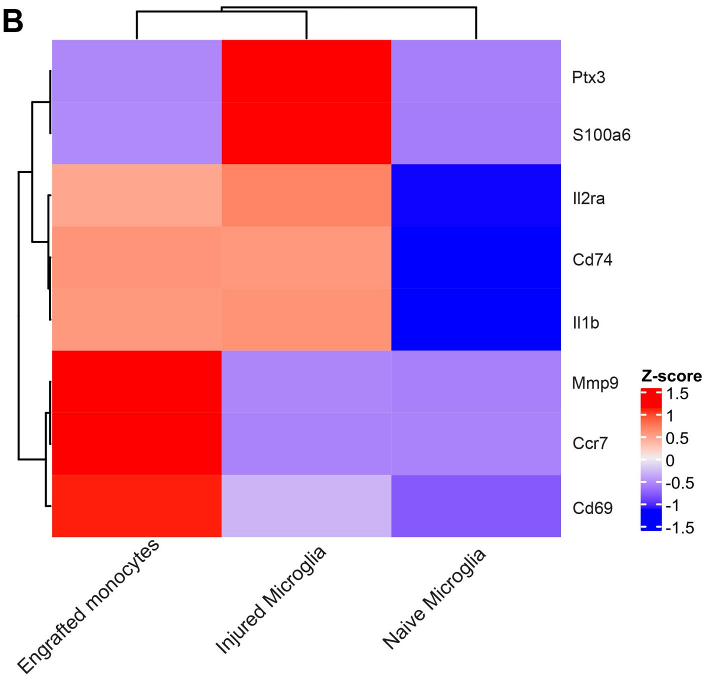


**C**

### NF-κB-associated genes

**Upregulated NF-κB-associated genes: injured vs naïve microglia**

| Gene | baseMean | log2FoldChange | padj |
|---|---|---|---|
| **S100a6** | 1045.012736 | 4.933703619 | 0.002430821 |
| **Cd74** | 478984.5346 | 7.805549164 | 2.43269E-10 |
| **Il2ra** | 2772.362103 | 13.04725963 | 3.17501E-06 |
| **Ptx3** | 65.01863895 | 8.46425414 | 0.002287336 |

**Upregulated NF-κB-associated genes: engrafted vs naïve microglia**

| Gene | baseMean | log2FoldChange | padj |
|---|---|---|---|
| **Mmp9** | 101.8758794 | 6.853878638 | 0.00073606 |
| **Cd74** | 474825.3241 | 7.902869561 | 7.39E-09 |
| **Il2ra** | 4747.756924 | 13.97011712 | 7.53E-14 |
| **Il1b** | 18229.04159 | 5.726035132 | 0.000129117 |
| **Cd69** | 7686.880832 | 4.515190221 | 0.011272649 |
| **Ccr7** | 33.46114622 | 8.398482004 | 0.011084859 |
